# Proteomic analysis of exosomes from lymphatic affluents reveals their implications in developing premetastatic niche in melanoma

**DOI:** 10.1101/2025.05.21.655399

**Authors:** Shankar Suman, Liyi Geng, Wendy K. Nevala, Xiaowei Zhao, Jaeyun Sung, James W. Jakub, Richard K. Kandasamy, Sarah A. McLaughlin, Akhilesh Pandey, Svetomir N. Markovic

## Abstract

Melanoma is an aggressive form of skin cancer that often spreads via lymphatic pathways to regional and distant sites. Melanoma-derived lymphatic exosomes play a crucial role in forming a tumor-supportive environment for metastasis, or premetastatic niche, within the first tumor draining lymph node, also known as the sentinel lymph node (SLN). Therefore, analyzing the proteomic content of tumor-draining lymphatic exosomes that deliver oncogenic molecules to the SLN is important in understanding the premetastatic niche. To reveal the proteomic landscape of lymphatic exosomes, we performed multidimension liquid chromatography-tandem mass spectrometry with multiplexing (18-samples) using tandem mass tag (TMT) labeling to profile the lymphatic exosomal proteomes obtained from afferent lymphatic channels leading to the SLN of patients with melanoma (n=6), control afferent lymphatic channels from prophylactic mastectomy (n=3) and non-cancer post-operative lymphatic fluid (n=9). Lymphatic fluid from postoperative lymphadenectomy drains served as another control to filter out non-melanoma alteration in lymph that may be related to the procedure of surgical resection and wound healing process. Our proteomic analysis identified 3929 proteins in the lymphatic exosomes, of which 968 were unique proteins absent from the current exosomal database. Interestingly, melanoma lymphatic exosomes possess distinctive proteomic cargo, which is significantly associated with cancer-associated and cellular structural remodeling pathways (FDR <0.05). Moreover, proinflammatory wound healing pathways are predominantly present in melanoma and postoperative lymph fluid compared to normal control afferent lymphatic channels. We identified a total of 17 uniquely modulated proteins in melanoma compared to control and postoperative lymph, which are critically involved in the process of melanoma tumorigenesis. At least ten upregulated proteins strongly correlate with each other at the gene expression levels in melanoma tumors compared to controls and may serve as a signature panel for melanoma (p = 2.13 x 10^-58^). In summary, this study represents the first comparative analysis of the lymphatic exosomal proteome and highlights distinct exosomal proteins that may support premetastatic niche formation in the SLN.

## Introduction

Melanoma is an aggressive form of skin cancer that can metastasize to regional lymph nodes and distant organs. A critical component of spread involves early metastases to the first tumor-draining lymph node (sentinel lymph node, SLN). We have previously described that this SLN is immunologically altered before the arrival of the first metastatic melanoma cells (pre-metastatic niche) (1). Melanoma-derived exosomes present in the lymphatic fluid draining from the primary tumor in the skin to the SLN contribute to the establishment of the immunotolerant microenvironment at the SLN, the premetastatic niche, fostering the colonization of melanoma cells (2). Earlier proteomic analysis of SLNs has shown that several upregulated proteins in SLNs can be indicative of the cancer state (3), which can be regulated by exosomes originating from primary tumor (4).

Lymphatic exosomes carry a rich source of tissue secretory factors and possess the ability to modulate systemic immune responses through the lymphatic route (5). However, following lymphatic drainage into the systemic circulation these exosomes are further diluted. Nonetheless, the proteomic cargo derived from plasma extracellular vesicles, which encompasses exosomes, has the ability to differentiate patients with melanoma from healthy subjects (6). However, the research focusing on exosomal proteomics within the lymphatic system in the context of melanoma remains limited, and no studies with comparator control lymph fluid have been conducted to date. During the progression of melanoma, lymphatic exosomes can penetrate the compromised macrophages in the subcapsular sinus of the lymph node, thereby facilitating cancer dissemination via lymphatic vessels (7). Furthermore, when endothelial cells receive melanoma exosomes, the expression of VE-Cadherin, EGFR and uPAR is enhanced to promote angiogenesis (8). Exosomal cargoes have been shown to reprogram stromal fibroblast to favor PMN development (9). Previous studies also show that exosomes derived from metastatic melanoma cells that propagate to the lung and brain have the capacity to trigger pro-inflammatory signaling pathways in lung fibroblasts and brain astrocytes, by recruiting immune cells through the expression of inflammation-activating factors, such as Hmgb1, Tslp, and Irf1 (10). Moreover, melanoma exosomes carry immunosuppressive properties as these exosomes directly dampen antitumor CD8+ T cell function (11).

There are limited studies on lymph proteome done to date. In 2004, the first lymph proteome was analyzed in normal Ovine lymph using SLEDI TOF-MS (12); however, human lymph proteomics remains sparse in the literature. A comparative study on human lymph and plasma was conducted using a bottom-up proteomics approach in 2014 by Clement et al. (13), which highlighted differences in composition between plasma and lymph. While several studies have probed blood-based exosomal proteomes across multiple cancers, including melanoma, there remains a significant gap in research regarding lymphatic exosomes. Our study is pioneering in its comparison of critical lymphatic exosomes with those previously documented from blood-derived exosomes, focusing on the differences between control and melanoma cases. It’s important to recognize that a lymph node acts as a site of “dynamic inflammation,” constantly undergoing changes in response to foreign threats. This process involves localized swelling and activation of immune cells, followed by a return to its normal size once the threat has been neutralized (14). To get a comprehensive understanding of changes in lymph nodes in melanoma induced by lymphatic exosomes, we conducted proteomic analysis of the lymphatic fluid samples obtained from axillary drains following lymphadenectomy. This approach was chosen because lymph node basin drains fluid serve as a unique cancer-free medium, enabling the separation of physiological inflammatory signals of wound healing from specific signals associated with melanoma, which are regulated by the lymphatics connected to melanoma tissues.

In our study, we focused on three distinct groups of samples for a thorough comparative analysis: (1) afferent lymphatic exosomes from non-tumor draining lymph nodes, which serve as a control; (2) afferent lymphatic exosomes obtained from SLNs of patients with melanoma: and (3), lymphatic exosomes from postoperative lymphadenectomy drains. All lymphatic exosomal protein samples were labeled with isobaric tandem mass tags for quantitative proteomics, which helps identify and quantify proteins within the lymphatic exosomal samples. Our approach aims to identify the melanoma exosomal cargoes that deliver oncogenic signals to lymph nodes, providing insights into melanoma-associated inflammation and potential therapeutic targets.

### Experimental Procedures

#### Informed consent, patient details and sample collection

We prospectively collected lymphatic fluid from afferent lymphatic channels leading from the primary melanoma to the SLN, as well as control lymphatic fluid obtained from afferent lymphatic channel from non-cancer surgeries (prophylactic mastectomy). These samples were collected by following the Declaration of Helsinki and approval by the Mayo Clinic Institutional Review Board (IRB: 10-000806). Lymphatic tracking using blue dye to the is a general clinical practice in melanoma operations to identify sentinel lymph node, which is the lymph node most likely to harbor metastasis. Using this method, the surgeon collected lymphatic channels leading from the primary cutaneous melanoma to the lymph node. The lymphatic channels were clipped at both ends to prevent lymph leakage before being submitted to the laboratory. For the control samples, the blue dye was injected into the breast in a standard subareolar/periareolar fashion to detect the blue afferent lymphatic channel draining into the axilla. These controls were selected from patients who were undergoing prophylactic mastectomies. We carefully collected lymph as a flow-through by rinsing the lymphatic channel with 1x PBS or RPMI solution using syringes. Furthermore, lymph fluid samples postoperative lymphadenectomy drain was collected under an approved clinical protocol (IRB# 08-004581). Informed consent was obtained from each participant for the prospective collection of biospecimens. The clinicopathological details of all biospecimens used are provided in supplementary table1. All lymphatic fluids were stored in the deep freezer until used in the experiment.

#### Exosome Isolation

Lymphatic samples were centrifuged at 500g for 10 minutes, followed by a second centrifugation at 2,000g for 20 minutes. From each sample, 1 ml was processed using a qEV1 70nm column (IZON) (Malvern, UK). The buffer configuration included 4 ml, with 0.7 ml collected per fraction, yielding a total of 6 fractions. The lymphatic exosomes were concentrated by passing them through a 30 kDa cellulose spin column (4 ml) at 3,000g for 15 minutes. The exosomes were then resuspended in 150 µl of PBS. The column was washed with 50 µl of 1% CHAPS, and both the wash and the resuspended EVs were mixed to a final volume of 200 µl. We measured the size and concentration of the samples using NanoSight NS30. The resulting concentrated samples were used for further downstream analysis, including transmission electron microscopy (TEM), immunoblotting, nanoparticle tracking analysis (NTA), and proteomic analyses.

#### Nanoparticle tracking analysis (NTA) of lymphatic exosomes

The particle size distribution and concentration of exosomal particles were analyzed using the NanoSight NS300 with NTA 3.4 software. Exosome samples were diluted 200-fold in PBS. Measurements were performed at a camera level of 14 and a syringe pump speed of 60. Particle tracking analysis was conducted with a detection threshold of 5. The exosomal particle concentration was reported per ml of sample after being measured in triplicate.

#### Transmission Electron Microscopy (TEM)

These vesicles were then adhered to carbon-coated 200 mesh copper grids. The grids were subsequently washed in 0.1 M phosphate buffer at pH 7.0 and fixed with a solution containing 4% paraformaldehyde and 1% glutaraldehyde. Finally, they were negatively stained with 1% phosphotungstic acid. Micrographs were further acquired using a JEOL1400 Plus transmission electron microscope (Peabody, MA).

#### Western Blotting

The exosomal lysates were first prepared with RIPA buffer, and protein concentration was measured by BCA protein quantitation kit (Thermo Fisher, Waltham, MA, USA) for western blotting. Equal protein concentration of blood plasma, lymph and lymphatic exosomes were loaded on a 4-15% gradient SDS-PAGE gel and run with an electrophoresis unit. The proteins on the gel were further transferred to PVDF membrane using Bio-Rad Wet/Tank Blotting Systems (Bio-Rad, Hercules, CA, USA) for 2 hours, which was confirmed by quick Ponceau S staining. Before probing the antibodies of exosomal markers, PVDF membranes were blocked with 5% non-fat milk in a 1× TBST solution. We incubated with primary antibodies against CD9 (cat #ab263019, abcam), Alix (cat #ab117600, abcam), Syntenin-1(cat #27964, cell signaling technology; CST), Albumin (cat #4929, CST), and calnexin (cat #ab22595, abcam) in blocking buffer under constant shaking for overnight at 4 °C. Furthermore, membranes were washed with 1X TBST (3x) and HRP labelled secondary antibodies against the host primary antibodies were added for 2 h at room temperature. Chemiluminescent substrates (Bio-Rad) were added to the membrane after removing unbound antibodies with 1x TBST (3x), and blots were imaged on X-ray films.

### TMT proteomics

#### Sample preparation and TMTpro labeling

To understand the remodeling of lymph nodes induced by melanoma exosome cargo, exosomes from these surgical biospecimens were further used for proteomics analysis. Isolated exosomes samples were processed using the S-Trap micro cartridges (ProtiFi.com) following the recommended protocol. Briefly, the samples were lyophilized and solubilized in 23ul of 5% SDS/50mM TEAB pH 8.5, reduced with 5mM TCEP at 50°C for 15 minutes, and alkylated with 10mM iodoacetamide for 20 minutes at room temperature before acidification with 2.5% phosphoric acid to create the protein suspension for loading into the S-trap cartridge. After several washes with 100mM TEAB / 90% MeOH, 20ul of (0.05ug/ul) Worthington trypsin/ 50mM TEAB pH 8.5 was added and incubated overnight at 37°C. The peptides are eluted with 0.2% formic acid and 0.1% TFA/ 50% acetonitrile and lyophilized. The peptides were solubilized in 100ul 300mM HEPES pH 8.5 buffer and labeled with 500ug TMTpro 18-plex reagents in 20 ul acetonitrile for 1 hour at room temperature and quenched with 5% hydroxylamine. Samples included afferent lymphatic channel of melanoma (n=6) and control (n=3) as well as postoperative lymph control (n=9). An 5ul aliquot from each label reaction was combined and analyzed by nanoLC-tandem mass spectrometry to confirm the labeling efficiency was >99%. The remainder of the reactions were pooled by normalizing with the reporter ion intensities to combine equal amounts of each sample. Excess TMT reagent was removed from the mixture using a Waters Sep-Pak C18 Plus long cartridge with 0.1% TFA /acetonitrile solvents, and the eluted the peptides were lyophilized.

#### Peptide fractionation with basic buffer HPLC

The dried TMT peptide mix was solubilized in 5% DMSO/5mM ammonium formate pH 8.5 and fractionated using basic pH reverse phase HPLC on a Thermo Ultimate 3000 RSLC HPLC system with a Waters XBridge peptide BEH C18, 3.5mm, 4.6mm column. The mobile phases were 5mM ammonium formate, pH 8.5 for the A solvent and 5mM ammonium formate, pH 8.5 / 90% acetonitrile/10% water for the B solvent. A flow rate of 0.5ml/minute was used, and gradient conditions were 5%B to 60%B over 60 minutes, to 80%B in 2minutes and hold for 5 minutes. A total of 96 fractions were collected over the 80minute method and were concatenated to 12 fractions and lyophilized prior to nanoLC-tandem orbitrap mass spectrometry analysis.

#### NanoLC-tandem Mass spectrometry Data Acquisition for total protein

The peptide fractions were analyzed by nanoLC-tandem mass spectrometry using a Thermo Scientific Orbitrap Eclipse mass spectrometer coupled to a Thermo Vanquish Neo UHPLC system with 0.1% formic acid in 98% water / 2% acetonitrile for the A solvent and 0.1% formic acid in 80% acetonitrile / 10% isopropanol / 10% water for the B solvent. Each fraction was solubilized in 0.1% formic acid and pumped onto a Halo C18 2.7µm EXP stem trap (Optimize Technologies, Oregon City, OR) with A solvent at a flow rate of 8µl / minute. The trap was placed in line with a Bruker PepSep C18 2.7um, 40cm x 100um column, and the peptides separated at a flowrate of 350nl/min with a gradient of 4%B to 40%B over 120minutes, then 40%B to 90%B over 12 minutes, with a 5-minute hold at 90%B for 5 minutes. Data were acquired using a real time search MS3 method for TMT quantitation. MS1 scans were performed in the Orbitrap with a mass range of 350-1500 m/z, AGC at 100%, 50ms max IT and resolution at 120,000 at 200 m/z. Ions were selected with an isolation width of 0.7 and fragmented by CID in the ion trap using the turbo scan rate and the MS2 spectra were searched in real time against a Swissprot human database with parameters set to allow 1 missed cleavage, static modifications of carbamidomethyl cysteine, TMT labeled lysine, TMT labeled peptide N-terminus and oxidized methionine as a variable modifications. Peptide spectral matches meeting minimum values of 1.2 for Xcorr, 0.1 for dCN and delta mass of 12 ppm or less, triggered an MS3 scan event for TMT reporter ion quantitation. Up to 10 fragment ions in the mass range 200-1500 from the matched spectra were isolated by synchronous precursor selection to generate the reporter ion MS3 spectra with HCD fragmentation at 55% NCE and orbitrap scanning from 100-500 m/z at 50k resolution. The MS3 AGC setting was 500% with the max ion injection time of 86ms. Dynamic exclusion was used to prevent ions selected for MS2 and any ions within an m/z of 7ppm from being selected for fragmentation for 25 seconds. The close out feature was implemented to restrict the MS3 trigger to a maximum of 6 peptide matches to a protein in each fraction run. The mass spectrometry proteomics data have been deposited to the ProteomeXchange Consortium via the PRIDE (15) partner repository with the dataset identifier PXD063898 and 10.6019/PXD063898’.

#### Protein Identification and Quantitation

The mass spectrometry raw data files were analyzed using Proteome Discoverer 3.0 (Thermo Scientific), set up for MS3 reporter ion quantification with TMT 6plex isobaric labels. The raw files were searched against a Swissprot human (2024_06) database using Sequest HT with parameters set for full trypsin specificity, allowing 2 missed cleavages and the variable modifications, oxidized Met and N-term protein acetylation. Fixed modifications were carbamidomethyl cysteine, TMT lysine and peptide n-terminal TMT. Mass tolerances were set at 10 ppm for precursor ions and 0.5 Dalton for MS2 fragment ions. Protein identifications were filtered at 1% FDR using the Percolator node. The MS3 TMT reporter ion channel intensities were reported with correction factors applied to PSMs and no filtering with isolation interference set at 100%. The matched proteins and raw reporter ion intensity values were exported to Excel and group comparisons were made with an in-house R-script.

### Statistical and Bioinformatics Analysis

To facilitate downstream analyses, TMT mass spectrometry proteomics data were pre-processed through missing feature exclusion, data normalization, imputation, and transformation. First, proteins with missing intensity values in more than 25% of the total samples were excluded. Next, sample-wise median centering was performed to normalize intensity values within each sample. Then, missing intensity values were imputed using half of the global minimum intensity value, which was identified as the smallest non-zero intensity value across all proteins and samples. Finally, all intensity values were log_2_-transformed. Statistical differences between groups were assessed using a two-sample *t*-test assuming unequal variances (Welch’s *t*-test), and differentially abundant proteins were identified based on [log_2_(mean fold-change)] ≥ 1.0 and *P*-value <0.05. Furthermore, volcano plots and other graphical representations were created using GraphPad Prism ver10 software. Lymphatic exosomal proteomes were compared to reported human exosome proteomic cargoes from the ExoCarta database (exocarta.org) and Vesiclepedia (microvesicles.org), and their comparison was made with Venn diagram analyses (VENNY 2.1). To gain insights into protein-protein interaction networks and conduct functional enrichment analysis of modulated exosomal cargo proteins, we utilized the STRING version 12.0 (https://string-db.org/) database. Additionally, we performed Gene Ontology (GO) analysis, Kyoto Encyclopedia of Genes and Genomes (KEGG) analysis, and other pathway enrichment analyses for the functional annotation of the proteomic cargoes using the DAVID and ShinyGO 0.82 webtools. We also used the TNMplotter web server (https://tnmplot.com/analysis) (16) to analyze gene expression, gene correlation (spearman correlation) and signature expression analyses for the identified altered proteomic signatures that uses RNAseq data from The Cancer Genome Atlas (TCGA)-Skin Cutaneous Melanoma (SKCM) datasets.

## Results

### 1. Isolation and characterization of lymphatic exosomes obtained from surgically dissected lymphatic channels

We established the collection of lymphatic fluid from the afferent lymphatic channels from control and melanoma cases, as shown in the methodology (Figure 1). Our study delineated the characterization of lymphatic fluid-derived exosomes and compared them with the source lymphatic fluid (Figure 2). Lymphatic exosomes were isolated from three different sources including afferent lymphatic channel of melanoma, afferent lymphatic channel of control (preventive mastectomy) and postoperative lymph fluid (collected post lymphadenectomy from breast surgery). Although the concentration of exosomal particles varied between samples, there were no significant differences of particle sizes among the three groups (Figure 2). Moreover, TEM analysis validates the NTA results we observed the exosomal particle size within 200nM which is key feature of small extracellular vesicle such as exosomes. We performed western blot to verify tetraspanins (CD9, CD63), ALIX (PDCD6IP), considered markers of exosomes and we also checked albumin (abundantly present in the plasma and lymph) and calnexin (an endoplasmic reticulum proteins which serve as negative marker of exosomes) (Figure 2D).

**Figure 1:**
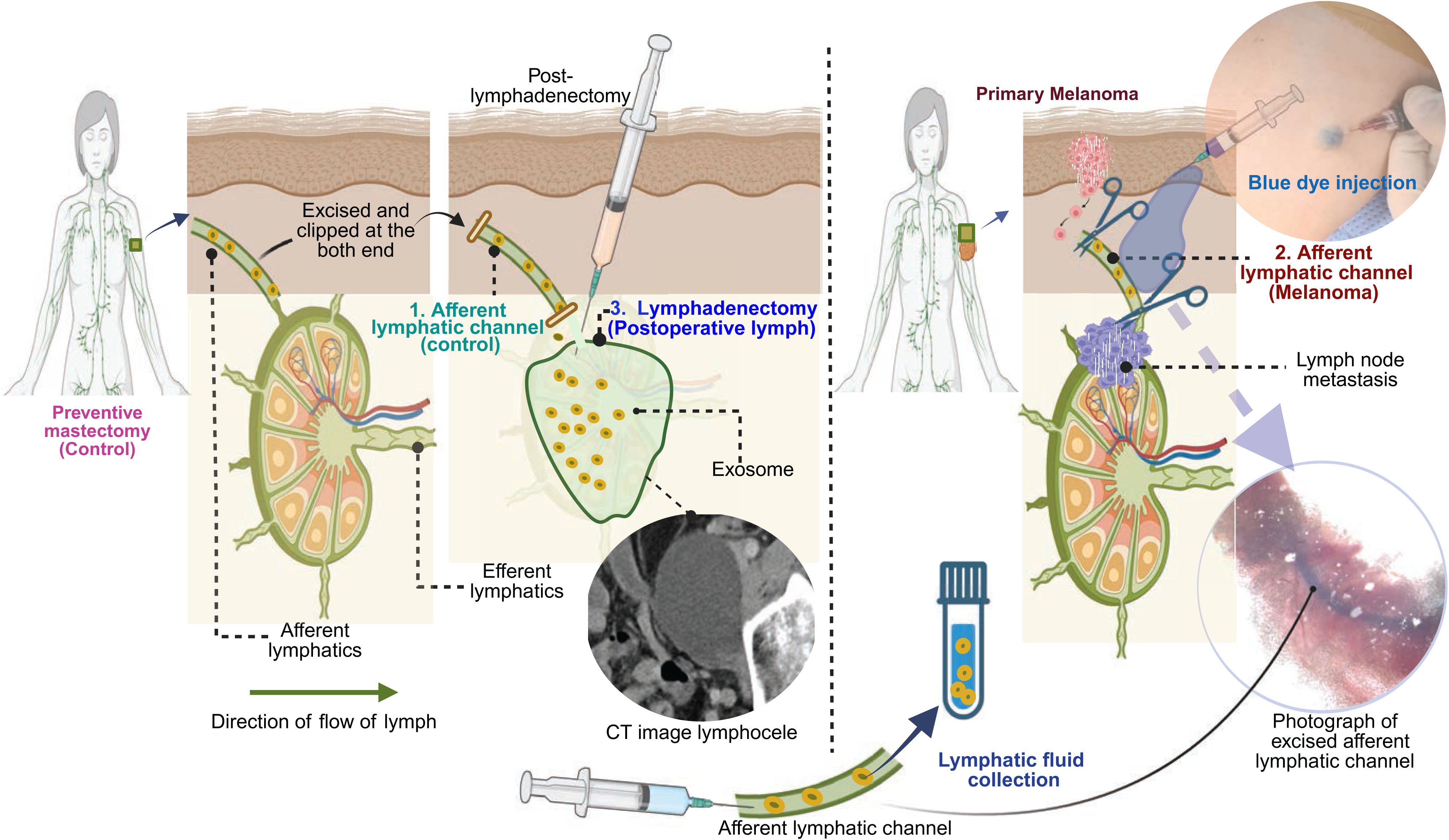
Collection of lymphatic fluid for exosome isolation from control cases (preventive mastectomy) and cutaneous melanoma cases. The afferent lymphatic channel which drains lymph from periphery tissue to sentinel lymph node was identified by surgeons using blue dye injection as part of their routine procedure. The excised intraoperative afferent lymphatic channel was clipped, and lymphatic fluid was collected by washing the channel with syringes from the following subjects 1. Individual undergoing preventive mastectomy (control), 2. melanoma patients undergoing lymphadenectomy for lymph node metastasis screening. Additionally, lymphatic fluid that drain post lymphadenectomy referred to as postoperative lymph fluid and categorized as 3. postoperative lymph. These three categories of samples were used isolation of lymphatic exosomes, and the proteomic analysis was performed.

**Figure 2:**
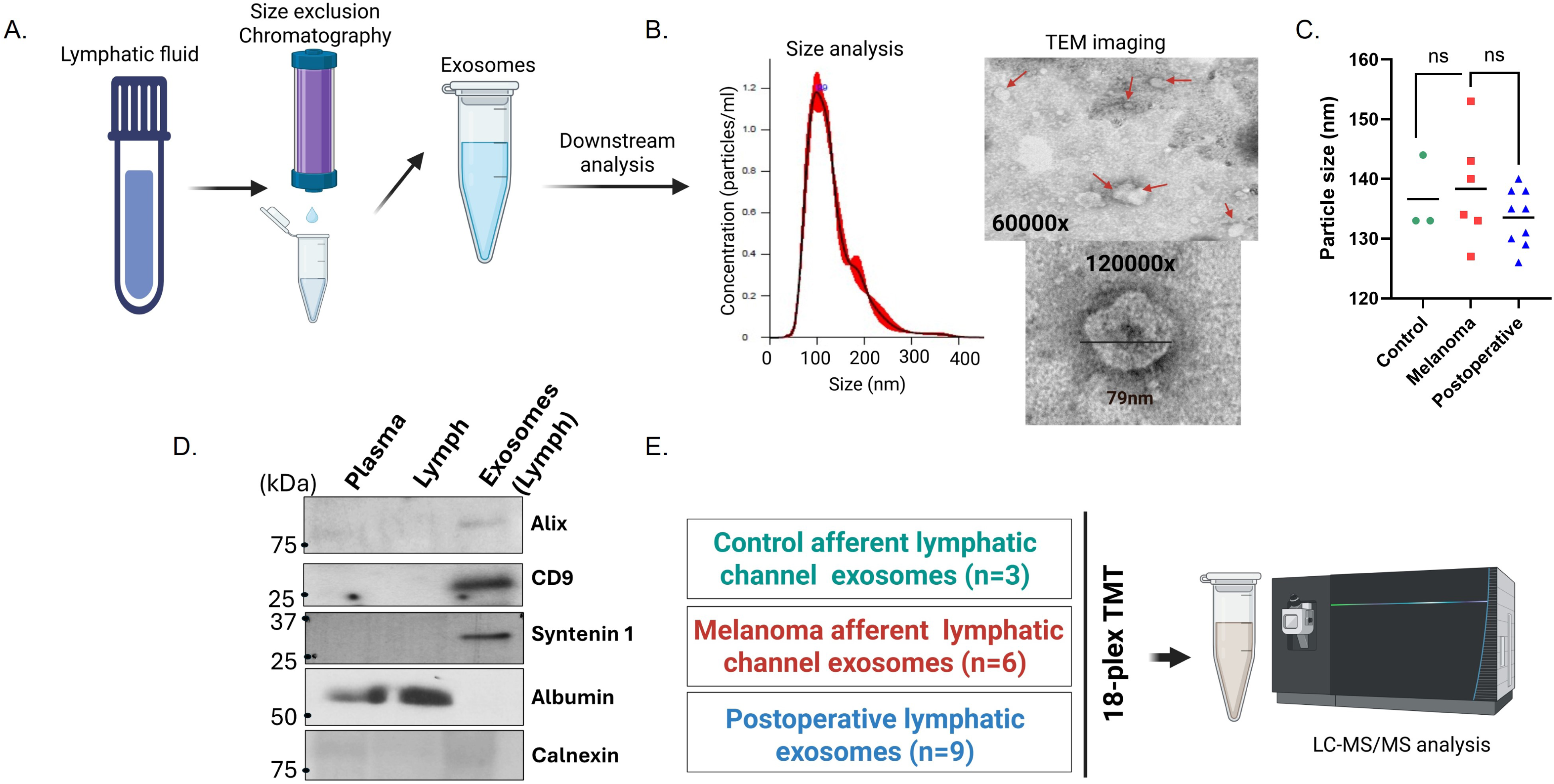
Experimental Workflow for Characterization of Lymphatic Exosomes and TMT Proteomics. **A.** Lymphatic fluid was used to isolate exosomal particles through size exclusion chromatography, followed by concentration using a cutoff column. Exosomes were characterized using nanoparticle tracking analysis (NTA) performed in triplicate. B. Transmission electron microscopy (TEM) analysis was conducted to examine the size and morphology of lymphatic exosomes. **C.** Dot plot indicates the average size of exosomes from each exosomal sample in control melanoma and postoperative lymph. **D.** Western blot analysis was performed to assess the presence of exosomal markers, including Alix, CD9, and syntenin-1, as well as negative makers albumin and calnexin in plasma, lymph, and exosomal particles. E. The TMT workflow involved labeling lymphatic exosomes collected from afferent channel control, melanoma and postoperative lymphatic fluid with 18 different TMT tags, which were further pooled, cleaned and then analyzed using mass spectrometry.

### 2. Proteome analysis of lymphatic exosomes identifies unique set of proteins

The dynamic nature of lymph nodes is largely influenced by lymphatic fluid, which comes from the peripheral tissue and is highly controlled as it enters the lymph node. Lymphatic exosomes can be unique in carrying cargo to regulate lymph node physiological function. Lymphatic exosomes are also considered efficient in delivering signals to the target organ by delivery into the systemic circulation, preparing lymph nodes for forthcoming threats (5). We analyzed the exosomal proteome from the lymphatic fluid from the surgically removed axillary lymph afferent channels, as well as the postoperative lymphatic fluid. The exosomes from the lymph identified 3,929 proteomic cargoes as identified using a stringent cutoff of the false discovery rate (FDR). Comparing lymphatic exosome proteome to all available cargoes of human exosomes available at ExoCarta, an exosome database collected from lymphatic channels, and Vesiclepedia, another compendium of extracellular vesicle’s cargoes, we identified 968 unique proteins in lymphatic exosomes (Supplementary file 1 & Figure 3A). Gene ontology analysis showed enrichment for proteins associated with organelle membranes and metabolic processes with oxidoreductase activity (Figure 3B). The most enriched proteins were COX6H2AX, NDUFA5, PLIN1, H2AC21, and ATP5F1B, each with coverage was over 50% (Figure 3C). Protein domain analysis revealed that the most significant proteins were linked with septin and septin-type guanine nucleotide-binding G-domains (Supplementary Table2). Septins have previously been reported as a cytoskeleton component that can self-assemble into a higher-order cytoskeleton and exhibit crucial functions in shaping cellular polarity, cell migration, vesicle transformation, and receptor signaling (17).

**Figure 3:**
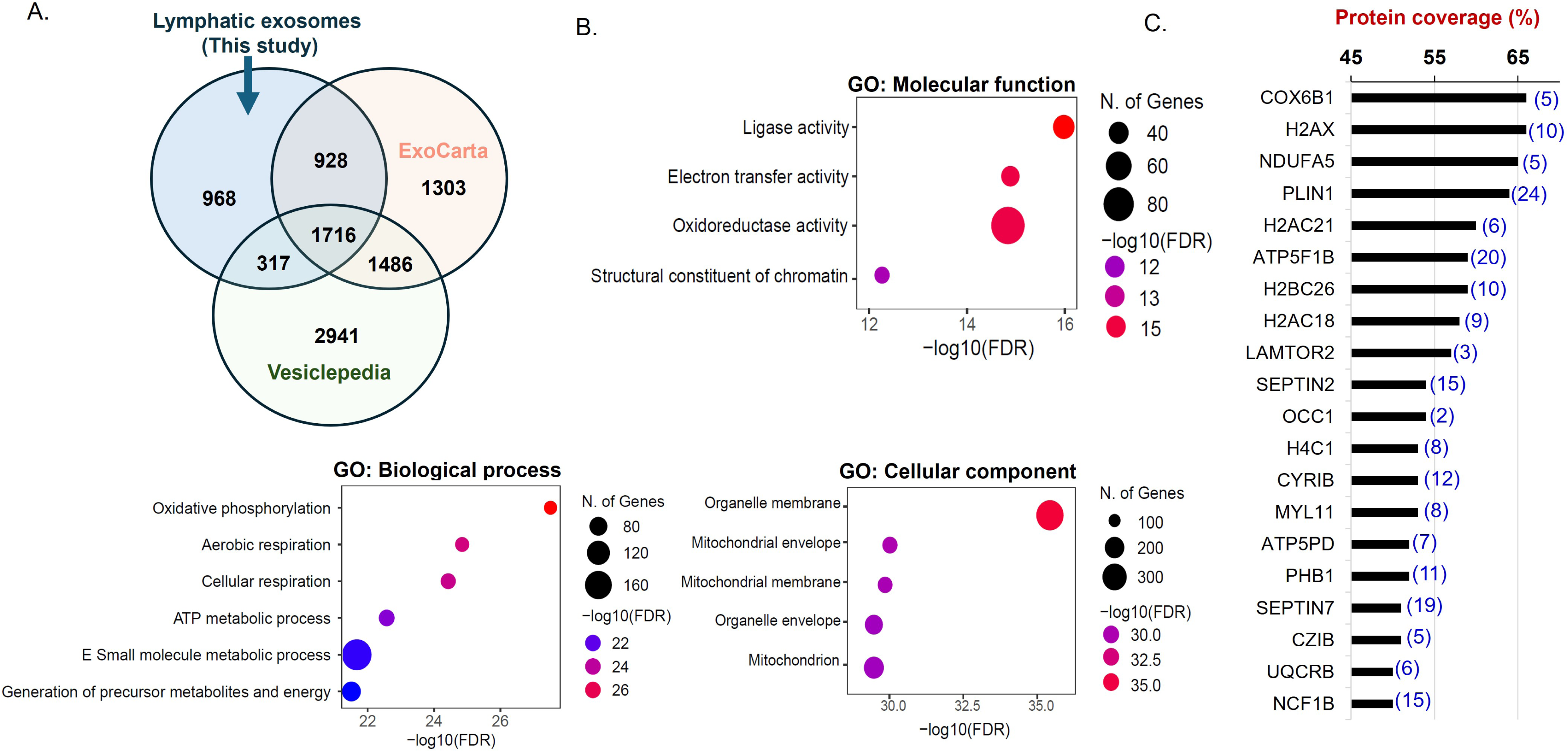
Lymphatic exosomes exhibit distinct proteomic cargo profiles that differ from those reported in datasets. **A.** Venn diagram displays the unique cargoes found in lymphatic exosomes, in comparison to the Human ExoCarta and Vesiclepedia datasets. **B.** The enrichment of protein domains and features of the unique proteins identified in lymphatic exosomes is depicted. **C.** A bar graph depicting protein coverage shows the top 20 lymphatic exosomal cargoes in lymph from this study, along with unique peptide matches (in blue).

Human melanoma cells primarily pass through the lymphatic system. Human lymph can be similar to blood plasma, but lymph can possess a higher concentration of tissue-secretory factors. Hence, we analyzed how the lymphatic exosomal proteome in melanoma compares to the exosomal profiles of human plasma and melanoma cells in the ExoCarta database. The reported exosomal proteins of human plasma show ∼38% overlap (93/244) with lymphatic exosomes; however, exosomes from reported human melanoma cells show ∼79% overlap (698/889) with lymphatic exosomes (Supplementary Figure 1). Moreover, 41 proteins are common in all, which fall under well-reported exosomal markers like tetraspanin (CD9, CD63, CD81), ESCRT-I proteins (TSG101), ALIX (PDCD6IP), heat shock proteins (HSP90AA1, HSP90B1), trafficking proteins (RAB5b), integrins (ITGB1), syntenin-1 (SDCBP), non-classical HLA class 1 family (HLA-E), and matched with top 10 reported exosomal proteins in the ExoCarta. Furthermore, western blot analysis confirmed the presence of exosomal proteins like ALIX, CD9, and syntenin-1 in the lymphatic exosomes (Figure 2D).

### 3. Altered proteomic map of exosomes from afferent lymphatic channel of melanoma and non-cancerous lymphatic channel control

Our studies analyze the proteome of lymphatic exosomes of melanoma versus non-control from the afferent lymphatic channels. In this setting, we identified 145 significantly altered proteins between control and melanoma lymphatic exosomes (p-value<0.05). Among them, 27 proteins were upregulated (>2-fold change, p-value <0.05), and 53 proteins were downregulated (>2-fold change, p-value <0.05) in the lymphatic exosomes of melanoma compared to control subjects (Figure 4A). Gene ontology analysis revealed differences in enriched proteins related to cellular components and post-translational modifications (PTMs) among the upregulated and downregulated protein cargoes in melanoma. Most of the downregulated proteins (34/53) were secreted proteins (FDR = 3.08 x 10^-17^), and 41/53 were glycoproteins (FDR = 1.54 x 10^-09^) in nature. However, no significant FDR was observed for cellular components or PTM-related proteins among the upregulated proteins in melanoma lymphatic exosomes (supplementary Table 3). Moreover, pathway analyses demonstrate that most upregulated proteins are associated with the enrichment of PD-L1 expression, and the PD-1 checkpoint pathway, as well as the Th1 and Th2 cell differentiation pathways (FDR <1 x 10^-2^) (Figure 4B). Notably, these pathways were primarily related to immune regulation during premetastatic niche formation (18). In contrast, the proteins that were downregulated are linked to pathways involved in ECM-receptor interactions, and the complement and coagulation cascades (FDR < 1 x 10^-4^) (Figure. 4B), all of which are highly relevant to the remodeling of the SLN (19).

**Figure 4:**
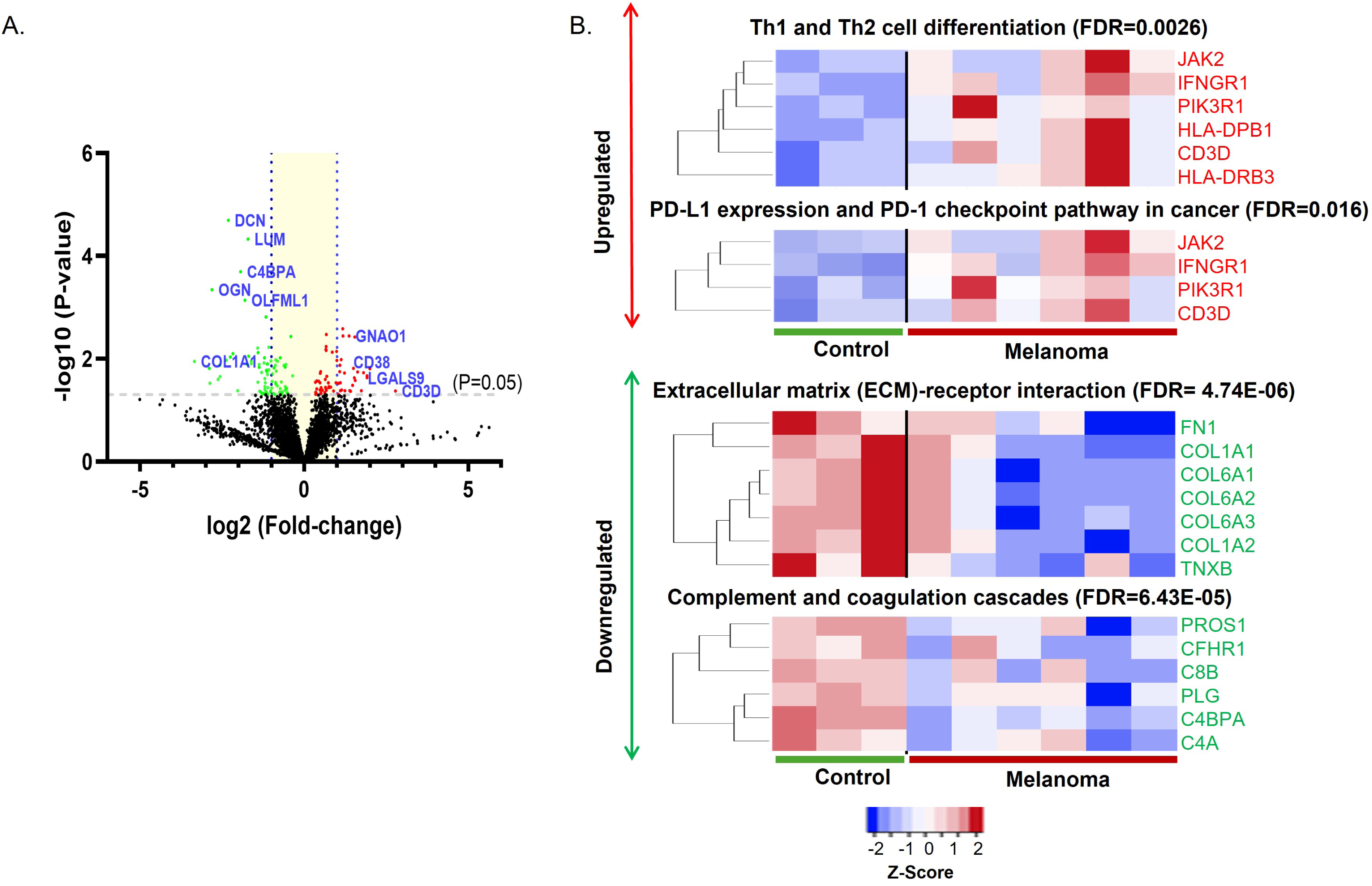
Melanoma lymphatic exosomes contain unique proteomic cargoes to establish oncogenic pathways in the SLN. **A.** The volcano plot illustrates the significant upregulation and downregulation of proteins in the exosomes derived from the afferent lymphatic channel of melanoma in comparison to the control group. **B.** KEGG pathway enrichment analysis reveals that modulated proteomic cargoes are associated with immunoregulation and cellular structural remodeling.

### 4. Comparative proteomics revealed proinflammatory and wound healing factors in melanoma lymphatic exosomes

We observed a significant alteration in the proteome of exosomes collected from the postoperative lymph in comparison to the control afferent channel (Supplementary Figure S2). One key reason for these changes is the presence of high tissue factors originating from resident immune cells surrounding the postoperative lymphs, which are primarily involved in wound healing and inflammatory responses. We compared the altered proteins in melanoma lymphatics to those in the control group and examined whether similar alterations were present in the postoperative lymph compared to the intraoperative control lymph. The common set of upregulated proteins identified in both groups highlight key proteins associated with wound healing and inflammation, which are also known features of the premetastatic niche. Among the differentially expressed proteins, we found those that are commonly upregulated in melanoma and postoperative lymph, as well as those uniquely upregulated and downregulated in melanoma group only (Figure 5A). More significantly, the PD-L1 pathway is common, which is key in both wound healing and tumor promotion (20). This process links IFNGR1 to JAK2, potentially culminating in the activation of PD-L1, which is a well-established immunosuppressive molecule (Supplementary Figure S3). The abundance of all these commonly upregulated proteins in postoperative lymph and melanoma samples is displayed in box plot graphs in Figure 5B, which corresponds to immune related pathways commonly perpetuate infection and wound healing process (21).

**Figure 5:**
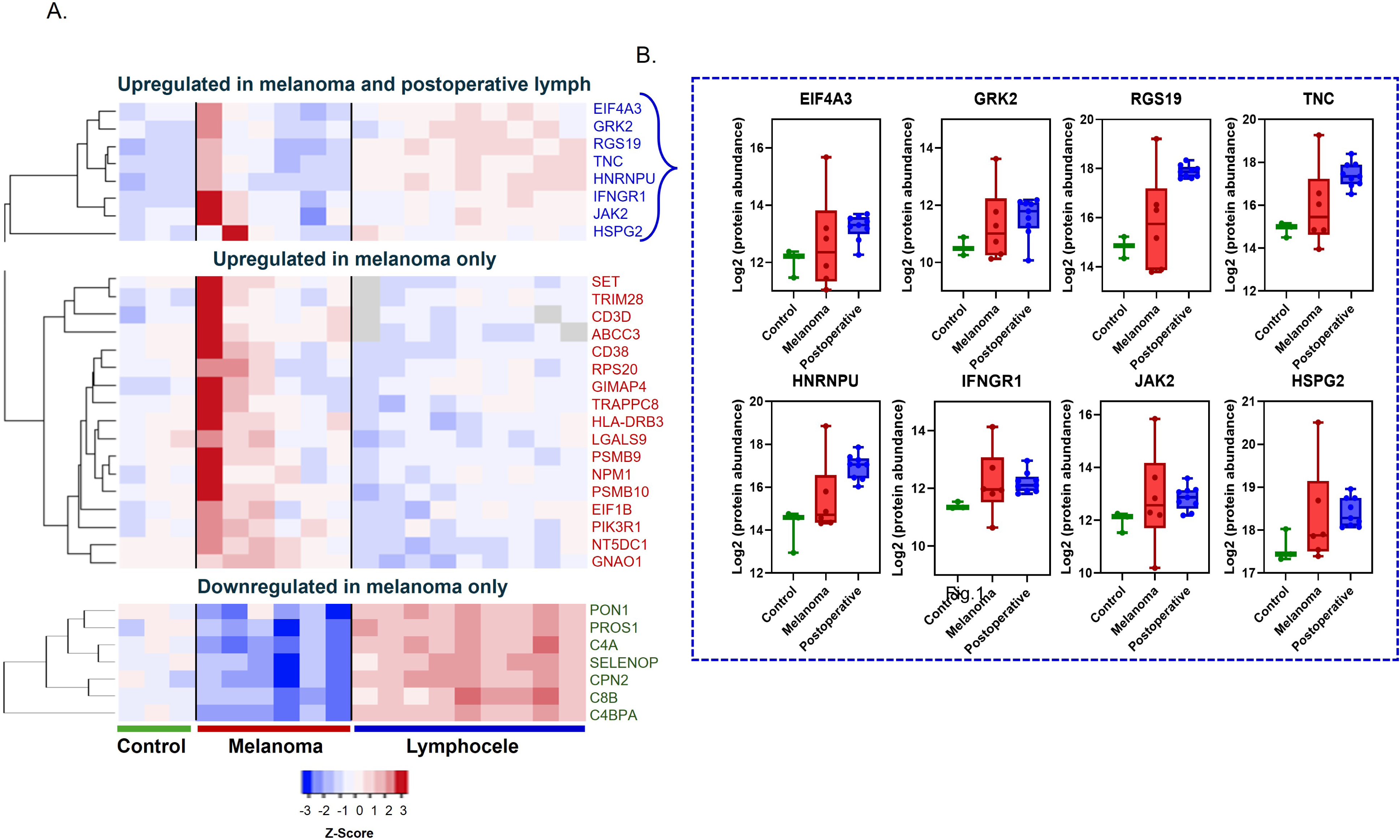
Proteomic analysis of control, melanoma and postoperative lymph exosomes demonstrates distinct proteomic cargoes of melanoma in wound healing process. **A.** Heatmap indicating exosomal proteomic differences among control, melanoma and postoperative lymph. **B.** Dot plots display common upregulated proteins of melanoma and postoperative lymph, compared to the control, are linked to proinflammatory and wound healing pathways.

### 5. Melanoma specific exosomal cargo proteins are involved in the melanoma metastasis

After analyzing all the upregulated and downregulated proteins in melanoma exosomes, we identified 17 proteins that were directly associated with melanoma metastasis based on the comparison of data from TCGA. This emphasizes the role of exosomes from afferent lymphatic channels in initiating tumorigenic pathways in recipient cells. The significance of these proteins was determined using skin cutaneous melanoma (SKCM) datasets using the TNM plotter webtool, which uses RNA sequencing data from normal tissues of non-cancer patients (474), tumor samples (103), and metastatic cases (368). Twelve uniquely upregulated proteomic cargoes in melanoma, compared to postoperative lymph or normal controls signifies their significance in the gene expression at the metastatic progression (Figure 6A). Additionally, the reduced gene expression of five downregulated proteins was significantly associated with melanoma metastasis (Figure 6B). These findings emphasize the potential of these candidates to be developed as biomarkers for the progression of melanoma metastasis.

**Figure 6:**
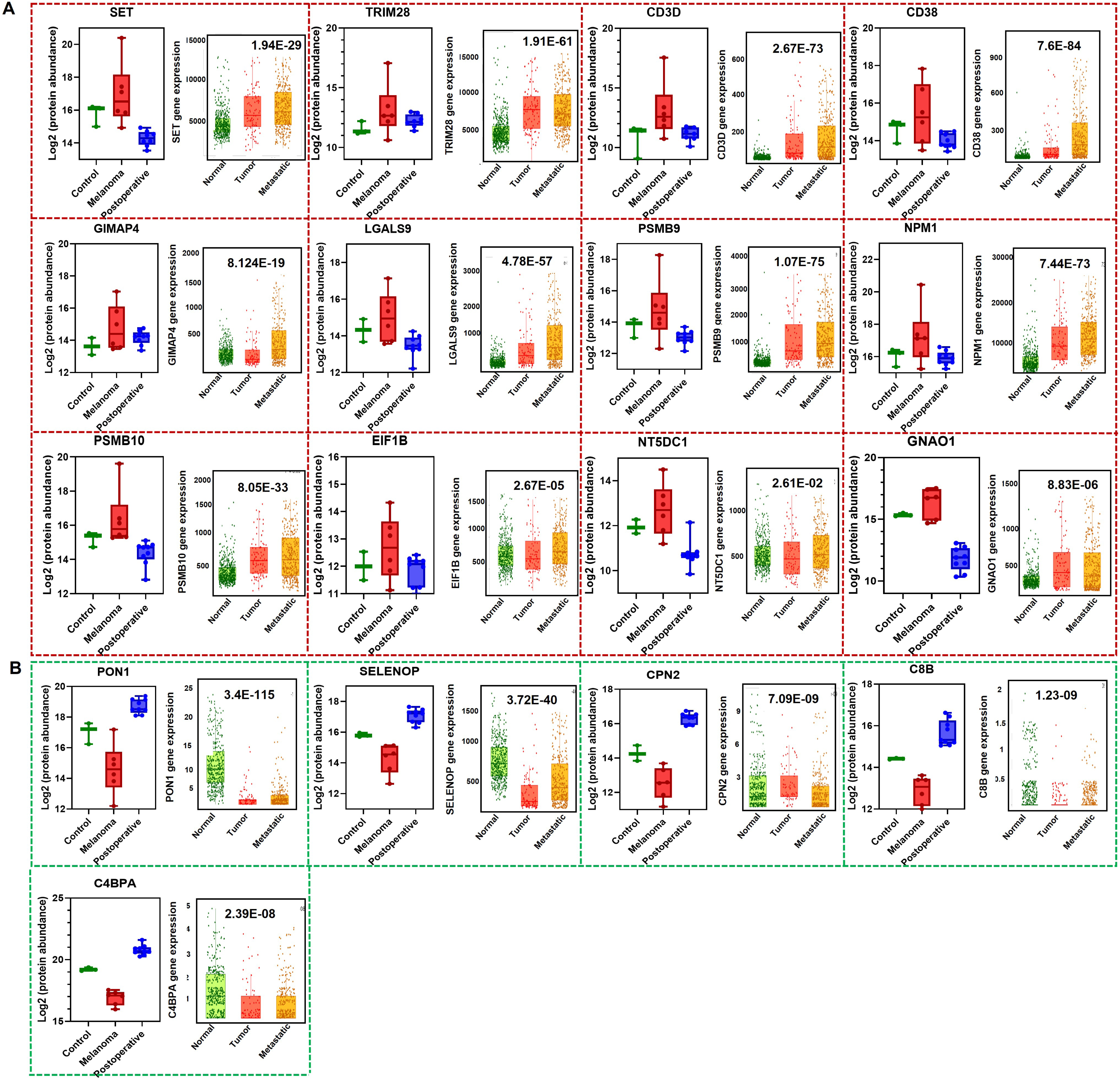
Primary melanoma communicates exosomal signals to facilitate tumorigenic pathways within the sentinel lymph node (SLN). Key melanoma-associated proteins have been categorized as **A.** upregulated and **B**. downregulated. The relative abundance of these proteins has been plotted for control, melanoma, and postoperative lymph samples. Furthermore, an analysis using TNMplotter indicates the direct involvement of the highlighted proteins in the melanoma metastasis. This analysis utilizes RNA sequencing data derived from 474 samples of normal tissues from non-cancer patients, 103 tumor samples, and 368 metastatic cases. These findings underscore the significance of unique proteomic cargoes within the context of melanoma.

### 6. Melanoma exosomes may hamper lymph node immunity through signalling cargoes to develop premetastatic niche

Primary melanoma-derived exosome-mediated immune suppression is well reported in SLN to support metastatic colonization (1, 2). Our proteomic studies revealed 79 proteins in the exosomes of melanoma lymphatics that may be involved in developing the pro-tumorigenic niche in the SLN. Those upregulated proteins link to several pathways in the SLN associated with inflammatory signals and wound healing in melanoma, as evidenced by postoperative lymph proteome alterations compared to controls (Figure 5B). Evaluating the differential expression levels of the upregulated proteins highlights the role of exosomal cargo proteins in myeloid cell dysfunction and CD8 T cell deactivation, among others supported by pathway analysis. In the proteomic network based on the MCL clustering method, the T-cell cell receptor (TCR) is predominant (Figure 7A). When we correlated the expression data at mRNA level, a stronger correlation of all 10 melanoma-specific cargoes was observed in the SKCM melanoma dataset compared to non-cancer normal tissue (Figure 7B). The signature panel analysis shows the higher significance of these upregulated, as well as downregulated (PON1, SELENOP, CPN2, C8B, C4BPA), candidates, indicating their relevance in melanoma metastasis (Figure 7C). Additionally, multiplex immunofluorescence (MxIF) analysis of selected candidate proteins like Tenascin C, Galectin-9, and CD38, revealed dominant staining against florescent labeled antibodies within SLN tissue, along with immune cells (CD45) and PDL1, which is a key immune checkpoint inhibitor molecule (Supplementary S4).

**Figure 7:**
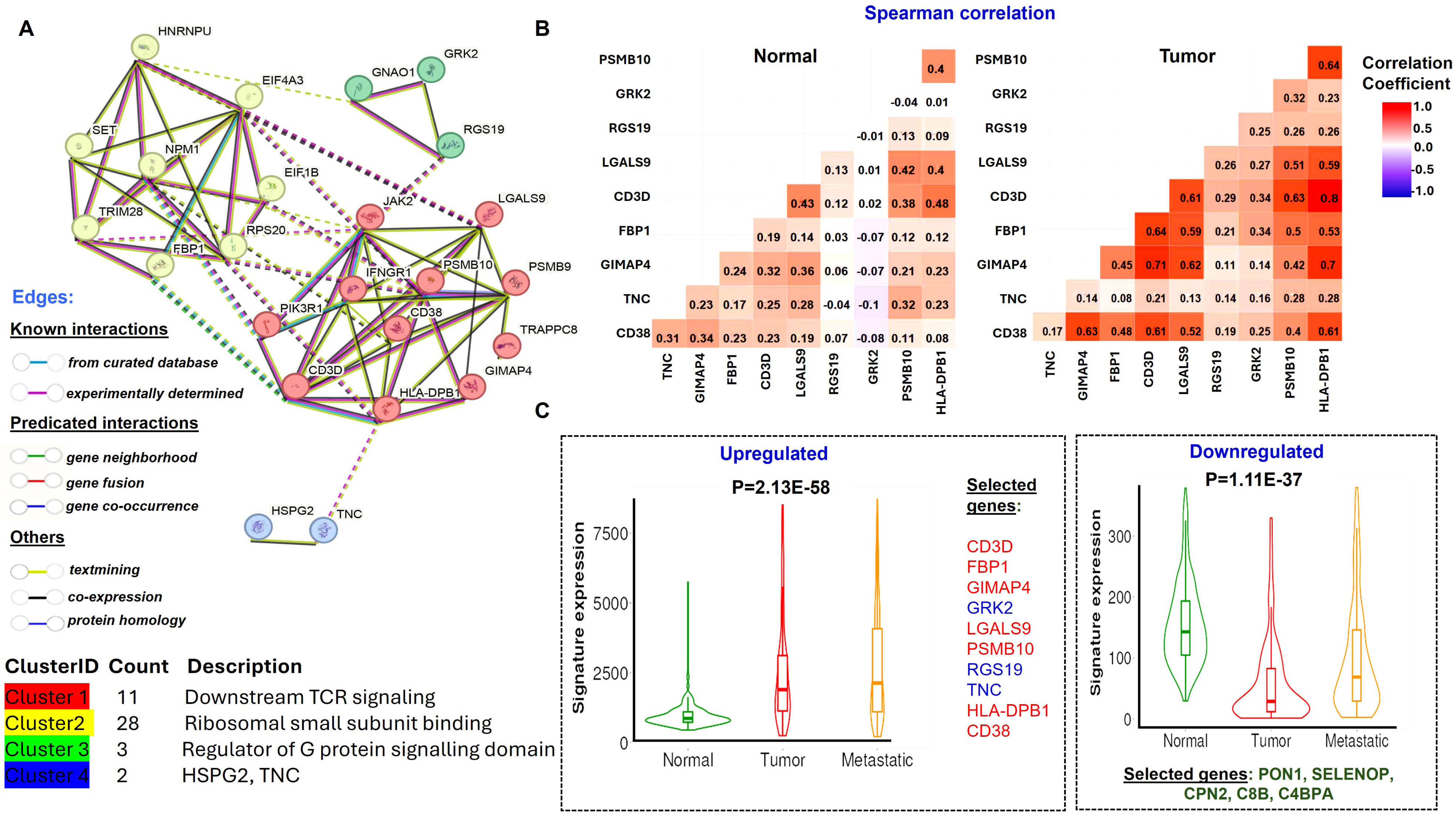
Significance of identified proteins derived from melanoma afferent lymphatic exosomes in melanoma tumorigenesis. **A.** The proteomic network of all upregulated proteins found in melanoma lymphatic exosomes, when compared to controls, reveals four significant protein network clusters identified using the MCL clustering method. **B.** A Spearman correlation matrix of the melanoma-associated upregulated proteins shows a higher correlation in melanoma tumor data compared to the normal dataset. **C.** Gene signature analysis using RNA-Seq data in relation to the SKCM dataset of the highlighted proteins includes a selection of upregulated and downregulated proteins in melanoma.

## Discussion

Tumor-derived exosomes influence the tumor microenvironment by facilitating intercellular communication, and they also have the potential to create a pro-tumorigenic niche. The cargoes within these exosomes are crucial for establishing the premetastatic niche, particularly in the lymph nodes where initial melanoma cell metastasis occurs. Our research provided insight into exosomes in the lymphatic fluid, which are crucial in setting up metastasis in the SLN. We utilized TMT proteomics to identify lymphatic exosomal proteomes and to quantitatively assess the difference in protein levels between melanoma-associated and non-cancer-associated lymphatic exosomes. Our pioneering study offers insights into deregulated proteomic cargoes into lymphatic exosomes in melanoma cases compared to controls with samples collected from afferent lymphatic channels. It is important to note that lymphatic fluid from the afferent channel is responsible for bringing unprocessed antigens in the fluid from tissue, which differs from that in efferent channel which carries processed fluid away from the lymph nodes. For this reason, afferent channel lymphatic fluid is crucial for our proteomic assessment. The study encompasses in-depth features of lymphatic exosomes collected from inoperative lymph of melanoma and control cases, as well as postoperative lymph resulting from lymphadenectomy. The study provided strong evidence of the dynamic nature of lymph through altering lymphatic exosomes, as postoperative lymph exosomal proteome composition significantly shifted from intraoperative lymphatic exosomes. Furthermore, postoperative lymph exosomes may serve as a critical control for wound healing mechanism implicated in lymphedema and lymphangiogenesis (22), key features of the premetastatic niche of SLN in melanoma (23). Hence, it helped us identify melanoma exosomal cargoes linked to inflammatory responses and other oncogenic activities.

Lymphatic exosomes are crucial mediators in the transport of signaling molecules from peripheral tissues to SLNs, where they play a key role in intercellular communication and trigger systemic immunity in response to various stimuli carried by the lymphatic fluid. However, the procurement of these lymphatic exosomes is fraught with challenges, largely due to their elusive nature and the intricate dynamics of lymphatic circulation. As a result, there has been a significant gap in the exploration of omics technologies aimed at studying the composition and functionality of exosomes derived from lymph of afferent lymphatic channels. Earlier studies from the exudative seroma obtained from lymphadenectomy of patients with stage III melanoma show enrichment in exosomal proteins resembling melanoma progression (24). Lymphatic exosomes from afferent channels were found to be unique when compared to all reported proteomic cargoes of ExoCarta and Vesiclepedia. Although lymphatic exosomes contain predominant established markers like tetraspanins, syntenin-1 and ESCRT proteins, 968 unique proteins were identified, primarily enriched with septin-type guanine nucleotide-binding domains, septins, tRNA synthetases, and linker histones, among others, that have not been reported earlier. Syntenin-1 has been identified as the most abundant protein and a potential universal biomarker (25), which was also detected in lymphatic exosomes. The KEGG pathway analysis of upregulated proteins in the melanoma lymphatic exosomes highlights the PD1 and PDL1 checkpoint pathway and Th1/Th2 cell differentiation pathway. However, the top hits among the downregulated proteins are associated with protein digestion and absorption as well as ECM (extracellular matrix) receptor interaction proteins (Figure 3B). In recent years PD1/PDL1 has been considered as an important target for immunotherapies against melanoma, as these proteins play a significant role in resisting T cells immune function (26, 27). Studies have also shown that Th1/Th2 imbalance is also a major issue in melanoma progression (28, 29). Our research validates that both of these crucial signals are directly regulated by melanoma exosomes in the lymphatic system, which contributes to the suppression of SLN immunity. Additionally, the structural remodeling of SLN particularly the expansion of the lymph nodes, is a key mechanism that prepares SLN for metastasis (30). Previous studies have provided evidence that lymphatic drainage is involved in this process (31). Our results indicate that several downregulated cargoes in the lymphatic channels of exosomes may be directly involved in biophysical remodeling.

Cancer and wound healing share common hallmarks, with compromised regulation of wound healing pathways facilitating cancer progression (32). This interplay involves the commandeering of wound healing “master regulators,” affecting treatment strategies for chronic wounds and cancer alike (33). The inflammatory response during wound healing can accelerate the growth of pre-neoplastic cells and promote tumor colonization (34, 35). Additionally, a postoperative lymph, which is a pocket of lymphatic fluid from disrupted drainage, can serve as a model in wound healing studies (36). Our studies have identified key pathways that mediate tumor-promoting functions, such as the INFR1-JAK2 and PDL1 pathways (37). Impaired interferon signaling is a common immune defect in many human cancers (38). PD-L1 is well reported to be induced by interferon gamma signaling in melanoma cells (39). These pathways serve as crucial signals in the wound healing process as well (40). The upregulated proteins indicate strong inflammatory signals that melanoma secretes to develop a premetastatic niche in SLN (Figure 5). We identified proteins that are uniquely overexpressed in melanoma lymphatics, which did not change in postoperative lymph compared to the control. From this analysis, 17 proteins were significantly associated with their roles in melanoma metastasis when compared with SKCM RNA sequencing data (Figure 6). The SKCM data also shows evidence of strong correlation of 10 candidates (CD3D, FBP1, GIMAP4, GRK2, LGALS9, PSMB10, RGS19, TNC, HLA-DPB1, CD38) in melanoma compared to non-cancer normal tissue. Interestingly most of these proteins are well known in multiple cancers including melanoma. For example, galectin-9 (LGALS9) expression correlates with poor prognosis in multiple human cancers (41). Our previous study shows that galectin-9 binds with myeloid cells and promotes a tumor-supportive microenvironment (42). CD38 is pro-tumoral factor involved in the stromal regulation for angiogenesis and metastasis (43) and is commonly reported in lung to brain metastasis (44). CD3D is well reported in gastric cancer and melanoma for its role in immune cell regulation(45, 46). Moreover, Tenascin-c (TNC) is reported as a critical regulator of melanoma progression (47, 48). Among the downregulated proteins, paraoxonase 1 (PON1) which protects against oxidative stress (49), and C4BPA, which inhibits cancer progression by promoting antitumor T cells (50), are notable. The decreased proteomic cargoes in melanoma highlight complement pathways, which are known to play a role in innate immunity and influence cancer progression (51).

In summary, our study provides the first comprehensive report on lymphatic exosomes and their proteomic alterations in the context of the afferent channel and postoperative lymph. By focusing on the analysis of lymphatic exosomal proteomics at the same site of collection within the afferent lymphatics, we were able to construct a detailed proteomic map that exhibits notable differences between melanoma cases and control subjects. Our results show that melanoma-associated exosomes carry key proteins that can potentially be involved in regulating SLN immunity through various signaling mechanisms that influence both innate and adaptive immunity. This analysis further revealed that the exosomes derived from melanoma contain distinct immune-related and structurally regulated proteins, which play a critical role in the formation of a premetastatic niche. These findings also suggest that melanoma exosomes are not merely byproducts of tumor progression but are actively involved in modulating the tumor microenvironment to facilitate metastasis. Functional studies on the effect of exosomes in recipient cells in SLN may contribute to the development of therapeutic targets to inhibit lymphatic metastasis in melanoma and potentially other cancers.

## Supporting information

suppple fig

## Acknowledgments

We acknowledge NIH to fund R01 grant (CA260259) to support this research. We also acknowledge assistance of the Mayo Clinic Proteomics Core, which is a shared resource of the Mayo Clinic Cancer Center (NCI P30 CA15083).

## Abbreviations

TMT: Tandem Mass Tag
PMN: Premetastatic niche
DAVID: database for annotation, visualization, and integrated discovery
DEP: differentially expressed protein
FC: fold change
NTA: nanoparticle tracking analysis
TBST: Tris-buffered saline containing 0.1% Tween-20
STRING: search tool for the retrieval of interacting genes/proteins
LC-MS/MS: liquid chromatography-tandem mass spectrometry
TEM: transmission electron microscopy

## References

1. Hood, J. L., San, R. S., and Wickline, S. A. (2011) Exosomes Released by Melanoma Cells Prepare Sentinel Lymph Nodes for Tumor Metastasis. Cancer Research 71, 3792–3801

2. Suman, S., and Markovic, S. N. (2023) Melanoma-derived mediators can foster the premetastatic niche: crossroad to lymphatic metastasis. Trends in Immunology 44, 724–743

3. Aboulouard, S., Wisztorski, M., Duhamel, M., Saudemont, P., Cardon, T., Narducci, F., Lemaire, A. S., Kobeissy, F., Leblanc, E., Fournier, I., and Salzet, M. (2021) In-depth proteomics analysis of sentinel lymph nodes from individuals with endometrial cancer. Cell Rep Med 2, 100318

4. Sun, B., Zhou, Y., Fang, Y., Li, Z., Gu, X., and Xiang, J. (2019) Colorectal cancer exosomes induce lymphatic network remodeling in lymph nodes. International Journal of Cancer 145, 1648–1659

5. Maillat, L., Potin, L., Kilarski, W. W., and Swartz, M. A. (2019) Abstract 4520: Lymphatic vessels regulate exosome trafficking from tumors. Cancer Research 79, 4520–4520

6. Bollard, S. M., Howard, J., Casalou, C., Kelly, B. S., O’Donnell, K., Fenn, G., O’Reilly, J., Milling, R., Shields, M., Wilson, M., Ajaykumar, A., Triana, K., Wynne, K., Tobin, D. J., Kelly, P. A., McCann, A., and Potter, S. M. (2024) Proteomic and metabolomic profiles of plasma-derived Extracellular Vesicles differentiate melanoma patients from healthy controls. Transl Oncol 50, 102152

7. Pucci, F., Garris, C., Lai, C. P., Newton, A., Pfirschke, C., Engblom, C., Alvarez, D., Sprachman, M., Evavold, C., Magnuson, A., von Andrian, U. H., Glatz, K., Breakefield, X. O., Mempel, T. R., Weissleder, R., and Pittet, M. J. (2016) SCS macrophages suppress melanoma by restricting tumor-derived vesicle-B cell interactions. Science 352, 242–246

8. Biagioni, A., Laurenzana, A., Menicacci, B., Peppicelli, S., Andreucci, E., Bianchini, F., Guasti, D., Paoli, P., Serratì, S., Mocali, A., Calorini, L., Del Rosso, M., Fibbi, G., Chillà, A., and Margheri, F. (2021) uPAR-expressing melanoma exosomes promote angiogenesis by VE-Cadherin, EGFR and uPAR overexpression and rise of ERK1,2 signaling in endothelial cells. Cell Mol Life Sci 78, 3057–3072

9. Shu, S., Yang, Y., Allen, C. L., Maguire, O., Minderman, H., Sen, A., Ciesielski, M. J., Collins, K. A., Bush, P. J., Singh, P., Wang, X., Morgan, M., Qu, J., Bankert, R. B., Whiteside, T. L., Wu, Y., and Ernstoff, M. S. (2018) Metabolic reprogramming of stromal fibroblasts by melanoma exosome microRNA favours a pre-metastatic microenvironment. Sci Rep 8, 12905

10. Gener Lahav, T., Adler, O., Zait, Y., Shani, O., Amer, M., Doron, H., Abramovitz, L., Yofe, I., Cohen, N., and Erez, N. (2019) Melanoma-derived extracellular vesicles instigate proinflammatory signaling in the metastatic microenvironment. Int J Cancer 145, 2521–2534

11. Chen, G., Huang, A. C., Zhang, W., Zhang, G., Wu, M., Xu, W., Yu, Z., Yang, J., Wang, B., Sun, H., Xia, H., Man, Q., Zhong, W., Antelo, L. F., Wu, B., Xiong, X., Liu, X., Guan, L., Li, T., Liu, S., Yang, R., Lu, Y., Dong, L., McGettigan, S., Somasundaram, R., Radhakrishnan, R., Mills, G., Lu, Y., Kim, J., Chen, Y. H., Dong, H., Zhao, Y., Karakousis, G. C., Mitchell, T. C., Schuchter, L. M., Herlyn, M., Wherry, E. J., Xu, X., and Guo, W. (2018) Exosomal PD-L1 contributes to immunosuppression and is associated with anti-PD-1 response. Nature 560, 382–386

12. Leak, L. V., Liotta, L. A., Krutzsch, H., Jones, M., Fusaro, V. A., Ross, S. J., Zhao, Y., and Petricoin, E. F., 3rd (2004) Proteomic analysis of lymph. Proteomics 4, 753–765

13. Clement, C. C., Aphkhazava, D., Nieves, E., Callaway, M., Olszewski, W., Rotzschke, O., and Santambrogio, L. (2013) Protein expression profiles of human lymph and plasma mapped by 2D-DIGE and 1D SDS-PAGE coupled with nanoLC-ESI-MS/MS bottom-up proteomics. J Proteomics 78, 172–187

14. Benahmed, F., Ely, S., and Lu, T. T. (2012) Lymph node vascular-stromal growth and function as a potential target for controlling immunity. Clinical Immunology 144, 109–116

15. Perez-Riverol, Y., Bandla, C., Kundu, D. J., Kamatchinathan, S., Bai, J., Hewapathirana, S., John, N. S., Prakash, A., Walzer, M., Wang, S., and Vizcaino, J. A. (2025) The PRIDE database at 20 years: 2025 update. Nucleic Acids Res 53, D543–D553

16. Bartha, Á., and Győrffy, B. (2021) TNMplot.com: A Web Tool for the Comparison of Gene Expression in Normal, Tumor and Metastatic Tissues. Int J Mol Sci 22

17. Ivanov, A. I., Le, H. T., Naydenov, N. G., and Rieder, F. (2021) Novel Functions of the Septin Cytoskeleton: Shaping Up Tissue Inflammation and Fibrosis. The American Journal of Pathology 191, 40–51

18. Wang, Y., Jia, J., Wang, F., Fang, Y., Yang, Y., Zhou, Q., Yuan, W., Gu, X., Hu, J., and Yang, S. (2024) Pre-metastatic niche: formation, characteristics and therapeutic implication. Signal Transduction and Targeted Therapy 9, 236

19. Pathania, S., Khan, M. I., Bandyopadhyay, S., Singh, S. S., Rani, K., Parashar, T. R., Jayaram, J., Mishra, P. R., Srivastava, A., Mathur, S., Hari, S., Vanamail, P., and Hariprasad, G. (2022) iTRAQ proteomics of sentinel lymph nodes for identification of extracellular matrix proteins to flag metastasis in early breast cancer. Sci Rep 12, 8625

20. Foster, D. S., Jones, R. E., Ransom, R. C., Longaker, M. T., and Norton, J. A. (2018) The evolving relationship of wound healing and tumor stroma. JCI Insight 3

21. Ivashkiv, L. B. (2018) IFNγ: signalling, epigenetics and roles in immunity, metabolism, disease and cancer immunotherapy. Nat Rev Immunol 18, 545–558

22. Jian, Y., Li, Y., Zhang, Y., Tang, M., Deng, M., Liu, C., Cheng, M., Xiao, S., Deng, C., and Wei, Z. (2024) Lymphangiogenesis: novel strategies to promote cutaneous wound healing. Burns & Trauma 12

23. García-Caballero, M., Van de Velde, M., Blacher, S., Lambert, V., Balsat, C., Erpicum, C., Durré, T., Kridelka, F., and Noel, A. (2017) Modeling pre-metastatic lymphvascular niche in the mouse ear sponge assay. Scientific Reports 7, 41494

24. García-Silva, S., Benito-Martín, A., Sánchez-Redondo, S., Hernández-Barranco, A., Ximénez-Embún, P., Nogués, L., Mazariegos, M. S., Brinkmann, K., Amor López, A., Meyer, L., Rodríguez, C., García-Martín, C., Boskovic, J., Letón, R., Montero, C., Robledo, M., Santambrogio, L., Sue Brady, M., Szumera-Ciećkiewicz, A., Kalinowska, I., Skog, J., Noerholm, M., Muñoz, J., Ortiz-Romero, P. L., Ruano, Y., Rodríguez-Peralto, J. L., Rutkowski, P., and Peinado, H. (2019) Use of extracellular vesicles from lymphatic drainage as surrogate markers of melanoma progression and BRAF (V600E) mutation. J Exp Med 216, 1061–1070

25. Kugeratski, F. G., Hodge, K., Lilla, S., McAndrews, K. M., Zhou, X., Hwang, R. F., Zanivan, S., and Kalluri, R. (2021) Quantitative proteomics identifies the core proteome of exosomes with syntenin-1 as the highest abundant protein and a putative universal biomarker. Nat Cell Biol 23, 631–641

26. Munhoz, R. R., and Postow, M. A. (2018) Clinical Development of PD-1 in Advanced Melanoma. Cancer J 24, 7–14

27. Gérard, A., Doyen, J., Cremoni, M., Bailly, L., Zorzi, K., Ruetsch-Chelli, C., Brglez, V., Picard-Gauci, A., Troin, L., Esnault, V. L. M., Passeron, T., Montaudié, H., and Seitz-Polski, B. (2021) Baseline and early functional immune response is associated with subsequent clinical outcomes of PD-1 inhibition therapy in metastatic melanoma patients. Journal for ImmunoTherapy of Cancer 9, e002512

28. Lauerova, L., Dusek, L., Simickova, M., Kocák, I., Vagundová, M., Zaloudík, J., and Kovarík, J. (2002) Malignant melanoma associates with Th1/Th2 imbalance that coincides with disease progression and immunotherapy response. Neoplasma 49, 159–166

29. Eftimie, R., Bramson, J. L., and Earn, D. J. D. (2010) Modeling anti-tumor Th1 and Th2 immunity in the rejection of melanoma. Journal of Theoretical Biology 265, 467–480

30. Suman, S., and Markovic, S. N. (2023) Melanoma-derived mediators can foster the premetastatic niche: crossroad to lymphatic metastasis. Trends Immunol 44, 724–743

31. Rohner, N. A., McClain, J., Tuell, S. L., Warner, A., Smith, B., Yun, Y., Mohan, A., Sushnitha, M., and Thomas, S. N. (2015) Lymph node biophysical remodeling is associated with melanoma lymphatic drainage. Faseb j 29, 4512–4522

32. MacCarthy-Morrogh, L., and Martin, P. (2020) The hallmarks of cancer are also the hallmarks of wound healing. Science Signaling 13, eaay8690

33. Sundaram, G. M., Quah, S., and Sampath, P. (2018) Cancer: the dark side of wound healing. The FEBS Journal 285, 4516–4534

34. Antonio, N., Bønnelykke-Behrndtz, M. L., Ward, L. C., Collin, J., Christensen, I. J., Steiniche, T., Schmidt, H., Feng, Y., and Martin, P. (2015) The wound inflammatory response exacerbates growth of pre-neoplastic cells and progression to cancer. The EMBO Journal 34, 2219–2236

35. Fukuda, K., Sugihara, E., Ohta, S., Izuhara, K., Amagai, M., and Saya, H. (2016) Injury promotes melanoma metastasis via wound healing process with periostin. Journal of Dermatological Science 84, e21

36. Lv, S., Wang, Q., Zhao, W., Han, L., Wang, Q., Batchu, N., Ulain, Q., Zou, J., Sun, C., Du, J., Song, Q., and Li, Q. (2017) A review of the postoperative lymphatic leakage. Oncotarget 8, 69062–69075

37. Holzgruber, J., Martins, C., Kulcsar, Z., Duplaine, A., Rasbach, E., Migayron, L., Singh, P., Statham, E., Landsberg, J., Boniface, K., Seneschal, J., Hoetzenecker, W., Berdan, E. L., Ho Sui, S., Ramsey, M. R., Barthel, S. R., and Schatton, T. (2024) Type I interferon signaling induces melanoma cell-intrinsic PD-1 and its inhibition antagonizes immune checkpoint blockade. Nature Communications 15, 7165

38. Critchley-Thorne, R. J., Simons, D. L., Yan, N., Miyahira, A. K., Dirbas, F. M., Johnson, D. L., Swetter, S. M., Carlson, R. W., Fisher, G. A., Koong, A., Holmes, S., and Lee, P. P. (2009) Impaired interferon signaling is a common immune defect in human cancer. Proc Natl Acad Sci U S A 106, 9010–9015

39. Garcia-Diaz, A., Shin, D. S., Moreno, B. H., Saco, J., Escuin-Ordinas, H., Rodriguez, G. A., Zaretsky, J. M., Sun, L., Hugo, W., Wang, X., Parisi, G., Saus, C. P., Torrejon, D. Y., Graeber, T. G., Comin-Anduix, B., Hu-Lieskovan, S., Damoiseaux, R., Lo, R. S., and Ribas, A. (2017) Interferon Receptor Signaling Pathways Regulating PD-L1 and PD-L2 Expression. Cell Reports 19, 1189–1201

40. Wang, X. H., Guo, W., Qiu, W., Ao, L. Q., Yao, M. W., Xing, W., Yu, Y., Chen, Q., Wu, X. F., Li, Z., Hu, X. T., and Xu, X. (2022) Fibroblast-like cells Promote Wound Healing via PD-L1-mediated Inflammation Resolution. Int J Biol Sci 18, 4388–4399

41. Yang, R., Sun, L., Li, C.-F., Wang, Y.-H., Yao, J., Li, H., Yan, M., Chang, W.-C., Hsu, J.-M., Cha, J.-H., Hsu, J. L., Chou, C.-W., Sun, X., Deng, Y., Chou, C.-K., Yu, D., and Hung, M.-C. (2021) Galectin-9 interacts with PD-1 and TIM-3 to regulate T cell death and is a target for cancer immunotherapy. Nature Communications 12, 832

42. Enninga, E. A. L., Chatzopoulos, K., Butterfield, J. T., Sutor, S. L., Leontovich, A. A., Nevala, W. K., Flotte, T. J., and Markovic, S. N. (2018) CD206-positive myeloid cells bind galectin-9 and promote a tumor-supportive microenvironment. J Pathol 245, 468–477

43. Ben Baruch, B., Mantsur, E., Franco-Barraza, J., Blacher, E., Cukierman, E., and Stein, R. (2020) CD38 in cancer-associated fibroblasts promotes pro-tumoral activity. Lab Invest 100, 1517–1531

44. Ben Baruch, B., Blacher, E., Mantsur, E., Schwartz, H., Vaknine, H., Erez, N., and Stein, R. (2018) Stromal CD38 regulates outgrowth of primary melanoma and generation of spontaneous metastasis. Oncotarget 9

45. Yuan, L., Xu, J., Shi, Y., Jin, Z., Bao, Z., Yu, P., Wang, Y., Xia, Y., Qin, J., Zhang, B., and Yao, Q. (2022) CD3D Is an Independent Prognostic Factor and Correlates With Immune Infiltration in Gastric Cancer. Front Oncol 12, 913670

46. Zhang, C., and Wu, S. (2023) Hypomethylation of CD3D promoter induces immune cell infiltration and supports malignant phenotypes in uveal melanoma. The FASEB Journal 37, e23128

47. Fukunaga-Kalabis, M., Martinez, G., Nguyen, T. K., Kim, D., Santiago-Walker, A., Roesch, A., and Herlyn, M. (2010) Tenascin-C promotes melanoma progression by maintaining the ABCB5-positive side population. Oncogene 29, 6115–6124

48. Fromme, J. E., Dummer, R., Mauch, C., and Zigrino, P. (2023) Tenascin C is a valuable marker for melanoma progression independent of mutational status and MAPK inhibitor therapy. Experimental Dermatology 32, 707–709

49. Medina-Díaz, I. M., Ponce-Ruíz, N., Rojas-García, A. E., Zambrano-Zargoza, J. F., Bernal-Hernández, Y. Y., González-Arias, C. A., Barrón-Vivanco, B. S., and Herrera-Moreno, J. F. (2022) The Relationship between Cancer and Paraoxonase 1. Antioxidants (Basel) 11

50. Sasaki, K., Takano, S., Tomizawa, S., Miyahara, Y., Furukawa, K., Takayashiki, T., Kuboki, S., Takada, M., and Ohtsuka, M. (2021) C4b-binding protein α-chain enhances antitumor immunity by facilitating the accumulation of tumor-infiltrating lymphocytes in the tumor microenvironment in pancreatic cancer. J Exp Clin Cancer Res 40, 212

51. Afshar-Kharghan, V. (2017) The role of the complement system in cancer. J Clin Invest 127, 780–789

