## Supplementary material for "Proteomic analysis of exosomes from lymphatic affluents reveals their implications in developing premetastatic niche in melanoma": supple table

Supplementary Table1: **Clinicopathological details of lymph samples**

|  | **Control** | **Melanoma** | **Postoperative** |
| --- | --- | --- | --- |
| No of subjects | 3 | 6 | 9 |
| Age | 58 ± 14.7 | 44 ± 18.9 | 58.5 ± 15.7 |
| Race | White:3 | White:6 | White: 8  African American: 1 |
| Surgery  : ALND  (Axillary lymph node dissection) | Mastectomy + ALND: 3 | ALND:6 | Mastectomy + ALND: 7  ALND only: 2 |
| SLN type | 0 | SLN (-): 7  SLN (+): 2 |  |

Supplementary Table 2: **INTERPRO: Protein Domain**

| **Sr** | **Term** | **Counts** | **P-Value** | **FDR** |
| --- | --- | --- | --- | --- |
| 1 | Septin | 9 | 5.00E-08 | 8.40E-05 |
| 2 | G_SEPTIN_dom | 9 | 1.00E-07 | 8.40E-05 |
| 3 | aa-tRNA-synth_I_CS | 9 | 1.00E-07 | 8.40E-05 |
| 4 | aa-tRNA-synth_II | 9 | 6.20E-07 | 3.80E-04 |
| 5 | aa-tRNA-synth_II/BPL/LPL | 10 | 2.20E-06 | 1.10E-03 |
| 6 | Rossmann-like_a/b/a_fold | 12 | 3.10E-06 | 1.30E-03 |
| 7 | tRNAsynth_Ia_anticodon-bd | 7 | 1.30E-05 | 4.60E-03 |
| 8 | WHEP-TRS_dom | 5 | 2.20E-05 | 6.40E-03 |
| 9 | H1/H5 | 6 | 2.30E-05 | 6.40E-03 |
| 10 | Plipid/glycerol_acylTrfase | 7 | 1.10E-04 | 2.80E-02 |

**Supplementary Table 3**: Cellular component and PTM analysis of modulated proteins in melanoma lymphatic exosomes

|  | **Count** | **FDR** | **Genes** |
| --- | --- | --- | --- |
| **Upregulated proteins: PTMs** | | | |
| KW-0597~Phosphoprotein | 20 | 0.210860454 | ABCC3, NPM1, RGS19, SET, IFNGR1, EIF4A3, TNC, HNRNPU, GIMAP4, TRAPPC8, PIK3R1, CD3D, PSMB10, GRK2, TRIM28, HLA-DPB1, RPS20, JAK2, EIF1B, FBP1 |
| KW-0007~Acetylation | 12 | 0.172255116 | NT5DC1, NPM1, SET, TRIM28, EIF4A3, HNRNPU, RPS20, PIK3R1, EIF1B, FBP1, PSMB10, PSMB9 |
| KW-0832~Ubl conjugation | 10 | 0.172255116 | NPM1, SET, TRIM28, IFNGR1, EIF4A3, HNRNPU, RPS20, PIK3R1, JAK2, HLA-DRB3 |
| KW-0013~ADP-ribosylation | 4 | 0.029787113 | GNAO1, NPM1, TRIM28, HNRNPU |
| **Upregulated proteins: Cellular components** | | | |
| GO:0016020~membrane | 18 | 0.004167623 | ABCC3, NPM1, RGS19, IFNGR1, EIF4A3, TNC, HNRNPU, PIK3R1, HSPG2, CD3D, GNAO1, NT5DC1, GRK2, HLA-DPB1, CD38, RPS20, JAK2, HLA-DRB3 |
| GO:0005886~plasma membrane | 12 | 0.446550097 | GNAO1, ABCC3, GRK2, RGS19, IFNGR1, HLA-DPB1, CD38, PIK3R1, JAK2, HLA-DRB3, CD3D, HSPG2 |
| KW-0472~Membrane | 12 | 1 | GNAO1, ABCC3, NT5DC1, GRK2, RGS19, IFNGR1, HLA-DPB1, HNRNPU, CD38, JAK2, HLA-DRB3, CD3D |
| CARBOHYD: N-linked (GlcNAc...) asparagine | 8 | 1 | ABCC3, IFNGR1, TNC, HLA-DPB1, CD38, HLA-DRB3, CD3D, HSPG2 |
| **Downregulated proteins: PTMs** | | | |
| KW-0325~Glycoprotein | 41 | 1.54E-09 | APCS, FKBP10, TNXB, COL14A1, PROS1, PON1, SLC43A1, C4BPA, PLG, PRELP, MAGT1, C8B, CPN2, C4A, GPNMB, OLFML1, CTSG, TMED4, LUM, CMA1, SERPINF1, FN1, KRT7, PCOLCE, PODN, SOD3, DCN, COL1A1, MFAP4, COL3A1, COL1A2, CFHR1, CEACAM6, COL6A2, CD109, COL6A1, SELENOP, OGN, COL6A3, FKBP9, FMOD |
| KW-1015~Disulfide bond | 31 | 2.20E-05 | APCS, TNXB, COL14A1, PROS1, PON1, C4BPA, PLG, PRELP, MAGT1, C8B, CPN2, C4A, OLFML1, CTSG, CPA3, LUM, CMA1, FN1, PCOLCE, LYZ, SOD3, DCN, COL1A1, COL3A1, COL1A2, CFHR1, CEACAM6, CD109, OGN, COL6A3, FMOD |
| KW-0379~Hydroxylation | 8 | 5.60E-05 | COL1A1, COL3A1, COL1A2, COL14A1, PROS1, COL6A2, COL6A1, COL6A3 |
| KW-0873~Pyrrolidone carboxylic acid | 6 | 1.14E-04 | COL1A1, COL1A2, LUM, SERPINF1, FN1, FMOD |
| **Downregulated proteins: Cellular components** | | | |
| KW-0964~Secreted | 34 | 3.08E-17 | APCS, TNXB, COL14A1, PROS1, PON1, C4BPA, PLG, PRELP, C8B, CPN2, C4A, OLFML1, CTSG, LUM, CMA1, SERPINF1, FN1, APOA4, PCOLCE, PODN, LYZ, SOD3, DCN, COL1A1, MFAP4, COL3A1, COL1A2, CFHR1, COL6A2, COL6A1, SELENOP, OGN, COL6A3, FMOD |
| KW-0272~Extracellular matrix | 16 | 1.13E-14 | TNXB, COL14A1, LUM, FN1, PRELP, PODN, DCN, COL1A1, MFAP4, COL3A1, COL1A2, COL6A2, OGN, COL6A1, COL6A3, FMOD |
| KW-0345~HDL | 2 | 0.37824533 | PON1, APOA4 |
| KW-0034~Amyloid | 2 | 0.37824533 | APCS, LYZ |
