## Supplementary material for "Proteomic analysis of exosomes from lymphatic affluents reveals their implications in developing premetastatic niche in melanoma": suppple fig

### Slide 1
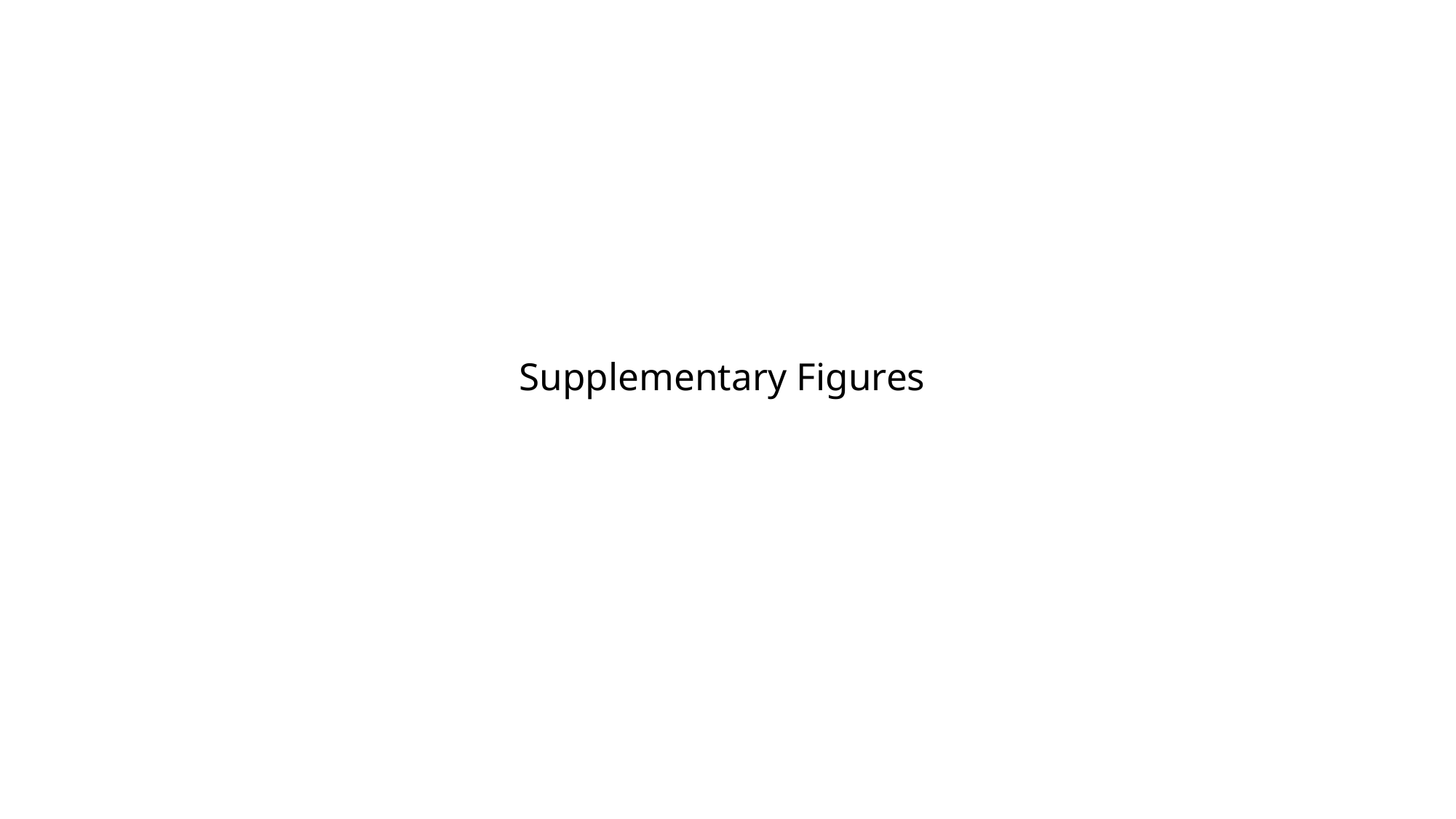

Supplementary Figures

### Slide 2
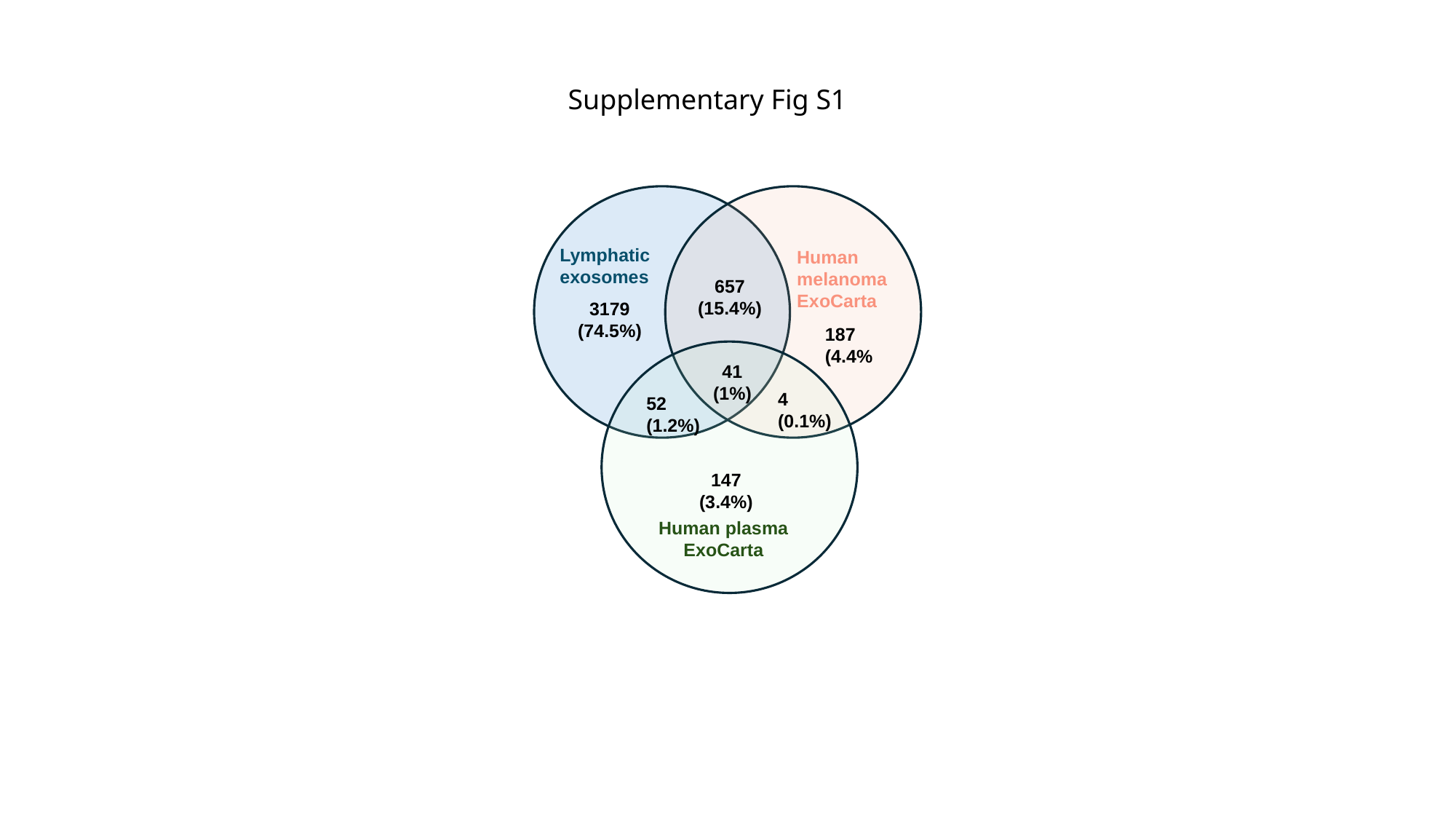

Supplementary Fig S1
Lymphatic exosomes
Human melanoma
ExoCarta
657
(15.4%)
3179
(74.5%)
187
(4.4%
41
(1%)
4
(0.1%)
52
(1.2%)
147
(3.4%)
Human plasma
ExoCarta

### Slide 3
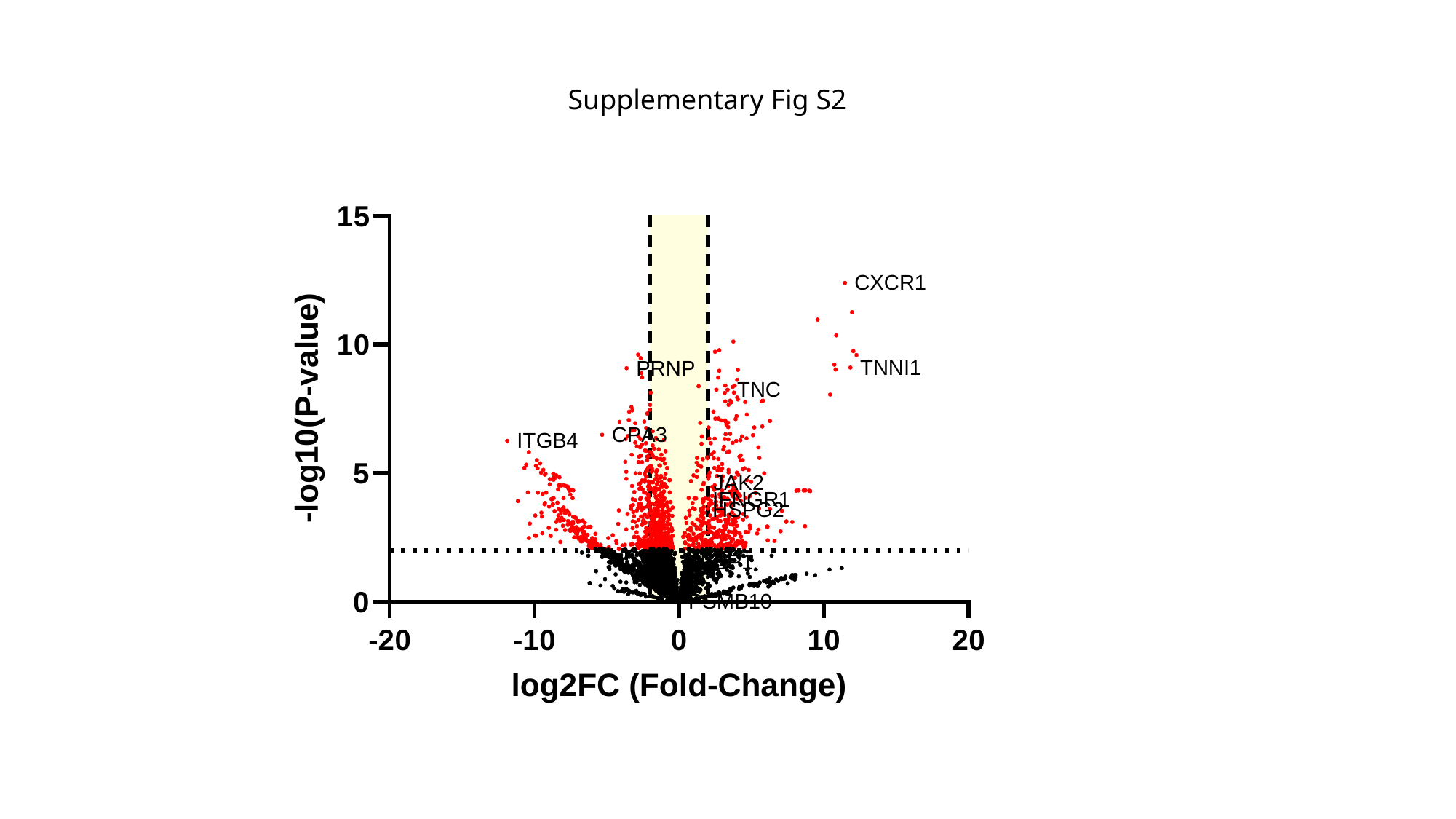

Supplementary Fig S2

### Slide 4
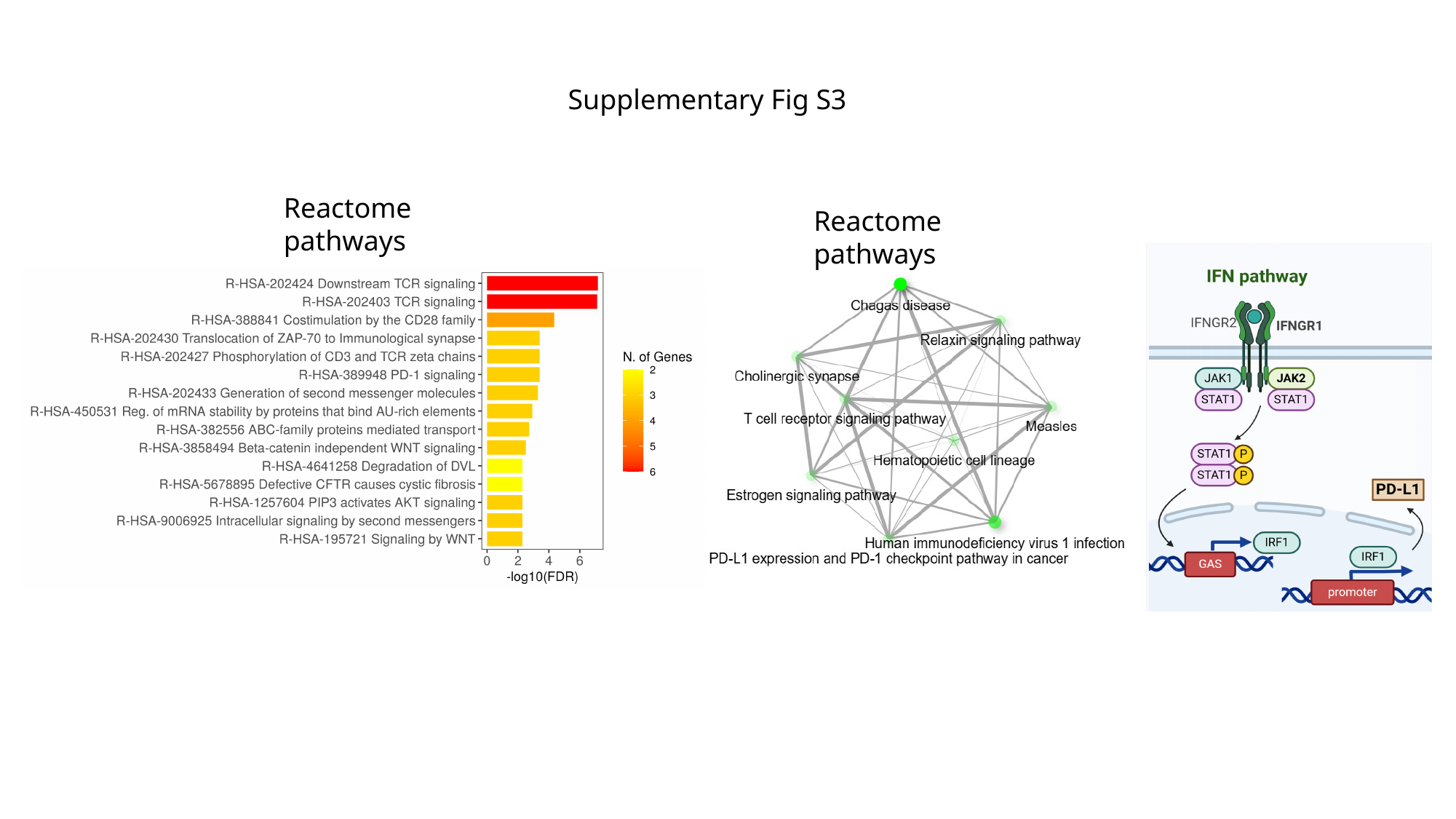

Supplementary Fig S3
Reactome pathways
Reactome pathways

### Slide 5
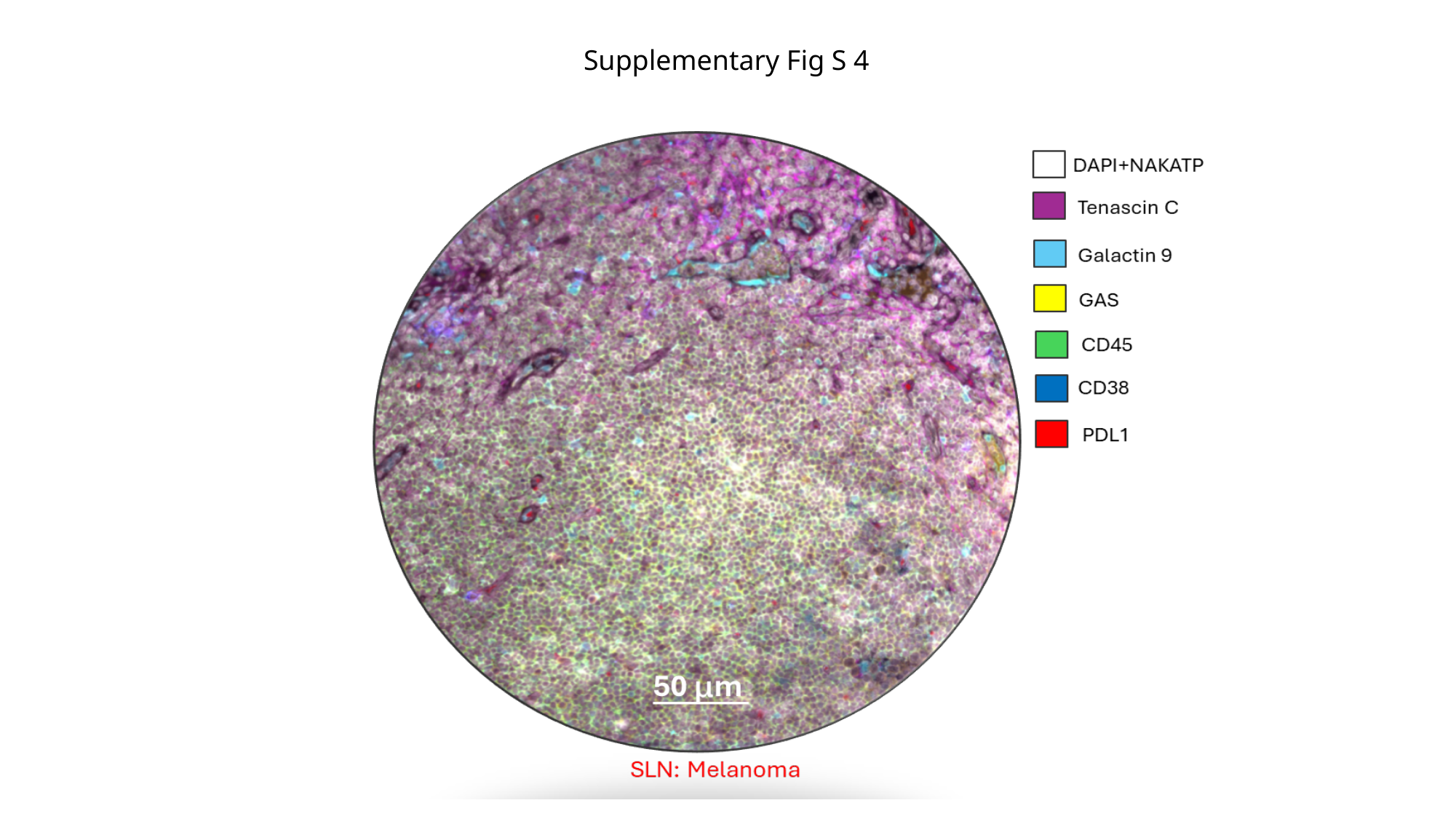

Supplementary Fig S 4
